# Tracking of *in vivo* O-GlcNAcylation in an Alzheimer’s Disease and Aging *C. elegans* Model

**DOI:** 10.1101/2025.07.16.659071

**Authors:** Fernando C. García Olivera

**Affiliations:** Department of Biochemistry, Faculty of Medicine and the Department of Biology, Faculty of Biology and Chemistry, Justus-Liebig University of Giessen, Giessen, Germany

**Keywords:** *C. elegans*, Aging, Alzheimer’s disease (AD), O-GlcNAc, CuAAC, JIPipe, visual programming analysis

## Abstract

We investigated O-linked β-N-acetylglucosamine (O-GlcNAc), a post-translational modification, in an *in vivo Caenorhabditis elegans* model of Alzheimer’s Disease and aging. Employing a chemoenzymatic labeling strategy combined with an automated image processing approach, we analyzed both post-hatching and adult stages of wild-type N2 and transgenic strain expressing human tau V337M under the *aex-3* promoter (*aex-3p*::tau(V337M)). Labeled O-GlcNAc proteins were visualized using fluorescence microscopy and quantified using a region-of-interest-based image analysis pipeline. Morphometric characterization revealed an age-dependent increase in O-GlcNAcylation in wild-type worms, while the AD model showed a progressive decline. In middle-aged transgenic nematodes, O-GlcNAc-labeled regions of interest shifted from anterograde to predominantly retrograde movement, suggesting that aging and neurodegeneration alter O-GlcNAc trafficking dynamics, potentially reflecting impaired synaptic support or enhanced clearance in *C. elegans*.

**Graphical Abstract:** 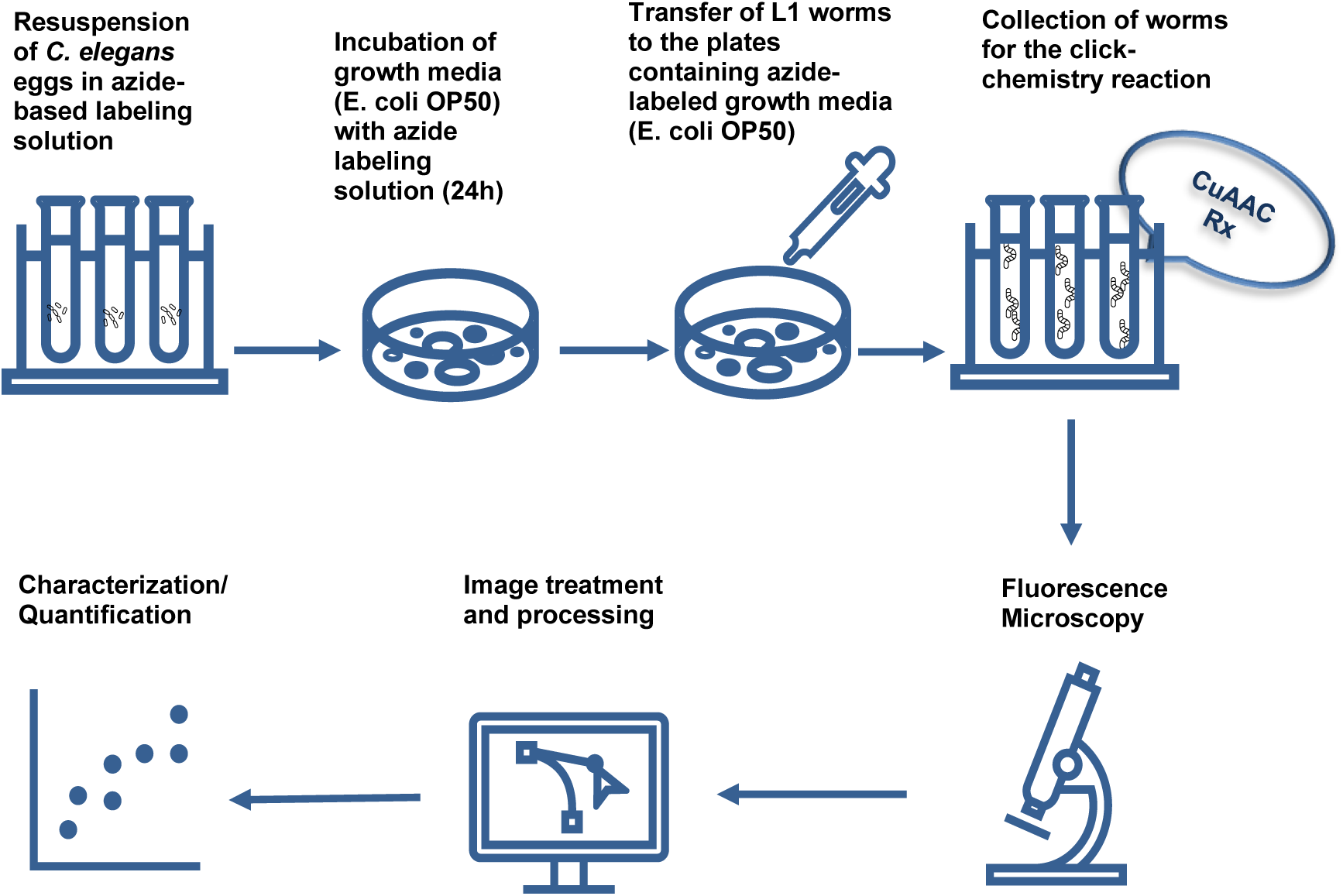

## Introduction

Alzheimer’s disease (AD) is the most prevalent form of dementia, characterized by progressive neurodegeneration and the accumulation of tau protein aggregates in the brain. Tauopathies, including AD, are defined by the abnormal hyperphosphorylation of tau, which impairs its microtubule-stabilizing function and disrupts cytoskeletal integrity (Frost, 2023). These changes contribute to synaptic dysfunction and neuronal loss, hallmarks of neurodegenerative decline.

Tau function is regulated by several post-translational modifications (PTMs), including phosphorylation and glycosylation, which influence its conformation, localization, and interactions (Ahmad *et al*., 2020). Among these, O-GlcNAcylation—the reversible attachment of a single N-acetylglucosamine moiety to serine or threonine residues—has emerged as a key modulator of neuronal protein dynamics. This modification is catalyzed by O-GlcNAc transferase (OGT) and removed by O-GlcNAcase (OGA). Its donor substrate, UDP-GlcNAc, is produced through the hexosamine biosynthetic pathway (HBP), linking O-GlcNAcylation to nutrient availability and cellular metabolic state (Cui *et al*., 2023).

O-GlcNAcylation regulates critical processes, including mitochondrial function, synaptic activity, and gene expression (Yutong *et al*., 2022; Xu *et al*., 2023; Varki *et al*., 2022). Notably, reduced O-GlcNAcylation correlates with tau hyperphosphorylation and impaired mitochondrial dynamics in AD, suggesting a protective role for this PTM (Pinho *et al*., 2019). Moreover, disruptions in O-GlcNAc homeostasis have been implicated in various diseases, including diabetes, cardiovascular disorders, and neurodegeneration (Ortega-Prieto *et al*., 2023; Costa *et al*., 2023). However, the *in vivo* dynamics of O-GlcNAcylation during aging and tau-mediated neurodegeneration remain insufficiently characterized.

Recent advances in metabolic chemical labeling have enabled visualization of glycosylation in live organisms. Azido-modified sugars can be incorporated into glycoconjugates and detected using click chemistry, allowing spatial and temporal tracking of glycosylated proteins (Ren *et al*., 2018; Battigelli *et al*., 2022). Despite their potential, these tools have been underutilized in models of neurodegeneration.

The nematode *Caenorhabditis elegans* is a genetically tractable model with a short lifespan, transparent body, and conserved neuronal pathways, making it well-suited for studying aging, metabolism, and neuronal function. O-GlcNAcylation is essential for normal *C. elegans* development and physiology (Wilson *et al*., 2023; Paschinger *et al*., 2023), and O-GlcNAc signaling in neurons has been linked to mitochondrial regulation and injury response (Taub *et al*., 2018).

In this exploratory study, we employed a transgenic *C. elegans* model expressing the human 4R1N tau isoform and the V337M AD mutation under the *aex-3* promoter (Pir *et al*., 2019; Alvarez *et al*., 2022). To visualize O-GlcNAcylated proteins *in vivo*, we combined metabolic labeling with click chemistry, followed by automated image analysis using JIPipe (Hoffmann *et al*., 2022; Gerst *et al*., 2023). This approach aimed to investigate preliminary patterns of *in vivo* O-GlcNAc dynamics during aging and in the context of AD to provide insights into this O-linked glycosylation.

## Materials and Methods

### Materials

Agar (Roth Cat. No. 8503); Peptone (Merck Cat. No. 91079-38-8); CaCl_2_ (Merck Cat. No. 102378); Cholesterol (Sigma-Aldrich Cat. No. C8503); CuSO_4_. 5 H_2_O (Sigma-Aldrich Cat. No. C8027); NaCl (Sigma-Aldrich Cat. No. S7653); K_2_HPO_4_ (Merck Cat. No. P290); KHCO_3_ (Sigma Aldrich Cat. No. 237205); KCl (Sigma-Aldrich Cat. No. 9541); KH_2_PO_4_ (Roth Cat. No. 3904.1); Na_2_HPO_4_ (Sigma-Aldrich Cat. No. S7907); MgSO_4_ (Roth Cat. No. 0261.1); NaOH (Sigma-Aldrich Cat. No. 1310-73-2); NaOCl (Roth Cat. No. 9062-3); EZClick O-GlcNAc Modified Glycoprotein Assay Kit™ (BioVision K714-100, Milpitas, CA, USA) containing: EZClick™ Wash Buffer (10X) cat. No. K714-100-1; Fixative solution Cat. No. K714-100-2; Permeabilization puffer (10X) Cat K714-100-3; EZClick^TM^ GlcAz Label (1000X) Cat. No. K714-100-4, Copper Reagent (100X) cat. No. K714-100-5; EZClick™ Fluorescent Alkyne (100X) Cat. No. K714-100-6; Reducing Agent (20X) cat. No. K714-100-7.

### Method Details

#### Worms Synchronization

Wild-type (WT) N2 (Bristol) and the *aex*-3p::tau(V337M) strains (University of Washington, Seattle, USA) were used for our experiments. The *aex-3* promoter drives the expression of the human tau V337M transgene predominantly in neurons and is commonly used in *C. elegans* for pan-neuronal expression, making it a suitable genetic tool for modeling tauopathy and investigating neuron-specific effects of post-translational modifications (Pir *et al*., 2019). A solid sterile Nematode Growth Medium (NGM) was prepared and sown with Escherichia Coli OP50 (Protein analytics, Giessen (HE), Germany) for feeding on 94 x16 mm plates (Greiner Bio-One, Frickenhausen, Germany) and was used to propagate *C. elegans* strains according to standard procedures (Stiernagle, 2006). All strains were grown at a temperature of 20°C. These plates were grown for 24h in an incubator (Incubator MAGV, Rabenau, Germany). Every two to three days, they were moved to a plate containing a new NGM medium. Adult worms were bleached using the alkaline hypochlorite method (Stiernagle, 2006) to prepare synchronized animals. In a nutshell, worms were removed from a p60 plate using M9 buffer (22 mM KH2PO4, 42 mM Na2HPO4, 86 mM NaCl and 1 mM MgSO4). After removing all the bacteria from the supernatant, they were lysed in a solution of alkaline hypochlorite for 1 minute and neutralized by using M9 medium; this process was monitored under a dissecting microscope and repeated six times until worms broke apart. Debris was washed with M9 by centrifugation at 2000 rpm for 3 min at 4°C (Hettich® Universal 32R Centrifuge, Tuttlingen, Germany) until the solution was cleared (eight wash cycles). Obtained eggs were then cleaned by centrifugation at 1800 rpm for 3 min with M9 medium at 4°C and allowed to nutate overnight before hatching in 2 mL of M9 for upkeeping in a climatic chamber (Liebherr®, Ochsenhausen, Germany) at 20°C.

#### O-GlcNAc Sites Labeling Protocol

To assess global levels of O-GlcNAcylated proteins, we employed the EZClick™ O-GlcNAc Modified Protein Detection Kit (BioVision), which utilizes a chemoenzymatic labeling strategy followed by click chemistry-based detection. This method is based on the metabolic incorporation of an azide-modified N-acetylglucosamine (GlcNAz) analog into O-GlcNAc residues, which are subsequently conjugated to a fluorescent reporter via copper(I)-catalyzed azide-alkyne cycloaddition (CuAAC). The approach offers high specificity and sensitivity for O-GlcNAc-modified proteins in whole-cell lysates and has been widely validated in the literature as a reliable strategy for profiling protein O-GlcNAcylation (Banahene *et al*., 2022) To obtain an azido label solution, an EZClick^TM^ GlcAz label solution was pipetted and mixed with 150 µL of E. coli (OP50) to a final 1XGlcAz concentrated culture and transferred onto freshly NGM p60 plates. The E. coli culture was completely spread on the plates and left to grow for 48h at 37°C.

150 µL aliquots of worm eggs of WT N2 (Bristol) and the *aex*-3p::tau (V337M) strains were pipetted in 15mL tubes containing 1mL of the azide (1xEZClick^TM^GlcAz) label solution.

Appropriate controls were included as follows:

##### Negative control

150 µl aliquots of worm eggs not exposed to the 1xEZClick^TM^GlcAz or treatment.

##### Positive Control

Worm eggs incubated with 1xGlcAz.

These were left nutating to hatch in the label-containing solution overnight at 20°C. Aliquots of hatchlings were seeded onto new NGM plates containing the azide-labeled OP50 culture (positive control) and onto a new NGM plate without the labeling reagent (negative control). Synchronized L1 *C. elegans (*young, day 1 post-hatched) of the corresponding strain were allowed to grow up to their “middle-aged” L4 stage (day 11 post-hatched) in the climatic chamber at 21°C. We utilized both the L1 and L4 larval stages to capture distinct aspects of neurodegenerative processes. L1-stage worms were used for early-stage exposure assays, taking advantage of their developmental sensitivity and ease of synchronization. L4-stage worms, with fully developed nervous systems, were employed to assess neuronal integrity and behavioral outcomes prior to adulthood. The use of both stages enables investigation of PTMs and neural vulnerability across key developmental points, as supported by prior studies on neurotoxicity and aging-related neurodegeneration (Taub *et al*., 2018; Wilson *et al*., 2023; Paschinger *et al*., 2023)

After the incubation with the labeling reagent, positive and negative control worms were removed and rinsed with fresh phosphate buffer saline (PBS) pH 7.4, discarding the supernatant. We cared that the suspended worms were pelletized at 500 x g for 5 min after the incubation procedure. PBS was gently removed with a pipette tip. Subsequently, 100 µL of the fixative solution was added to the worms and incubated for 15 min at room temperature (RT), protected from light. Fixative was removed, and the worms were washed twice with 1X wash buffer. The wash was removed, and 1X permeabilization buffer was then added to the worms, which were incubated for 10 min at RT. Permeabilization buffer was finally removed, and 1xEZClick^TM^ Reaction cocktail was prepared as follows:

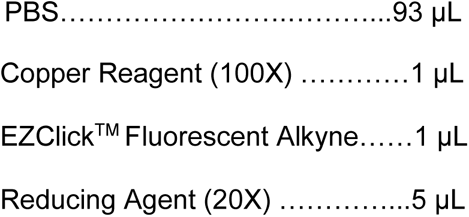

We added 100 µL of 1xPBS to resuspend the negative control worms. Separately, 100µL of 1 xEZClick^TM^ Reaction cocktail was added to L1 and L4 stages of N2 and *aex*-3p::tau (V337M) strains and incubated for 30 min at RT, protected from light. The reaction cocktail was removed and washed three times with the wash buffer. After the third washing cycle, the wash buffer was removed, and the worm samples were suspended in 100 µL PBS.

The negative and positive control samples were monitored before the copper-mediated click reaction under the fluorescent microscope. No signals were emitted from the worms not exposed to the alkyne dye labeling solution (Fig. 1A).

**Figure 1.**
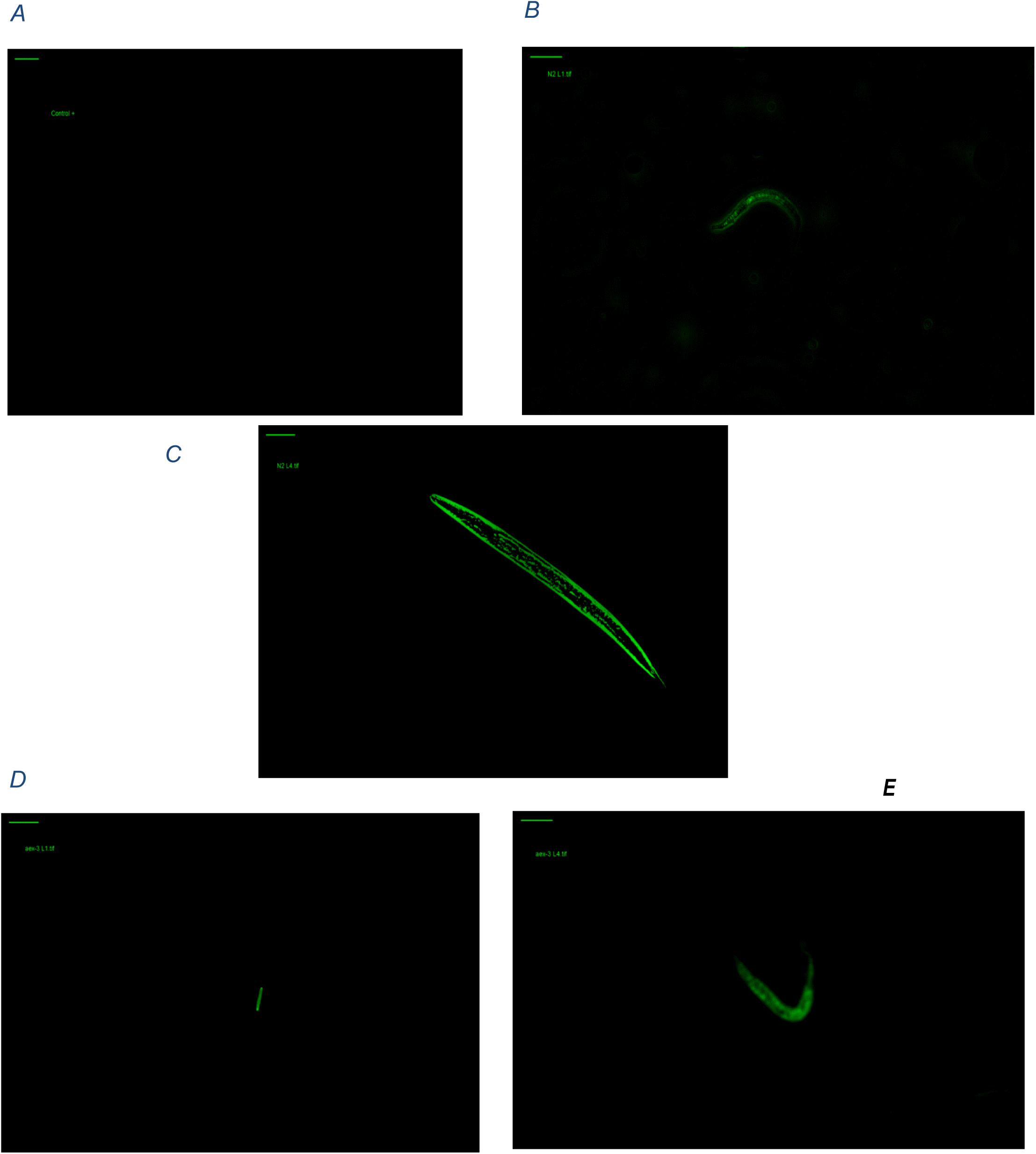
High-resolution fluorescence images of *in vivo* localization of O-GlcNAc sites in larval and adult C. elegans. (A) Represents worms only incubated with the azide sugar (control+GlcNAz) did not exhibit fluorescence. (B) N2 L1, (C) 11-day-old N2 L4, (D) *aex-3*p::tau (337M) L1 and (E) 11-day-old *aex-3*p::tau (337M) strains were incubated with GlcNAz and reacted with an alkyne-containing fluorescent dye under a copper-catalyzed cycloaddition. (B-E) Micrographs were converted to .tiff files and observed fluorescence was measured using a JIPipe workflow. Scale bars for all panels adjusted to 100µm.

### Microscopy Settings

We selected copper-catalyzed azide–alkyne cycloaddition (CuAAC) for O-GlcNAc detection due to its high specificity, cellular permeability, and compatibility with live-organism labeling, which are critical advantages for dynamic, *in vivo* studies in *C. elegans*. Although we did not perform additional biochemical confirmation (e.g., Western blotting or LC-MS/MS), the CuAAC-based metabolic labeling strategy has been extensively validated for glycoprotein visualization in other model systems (Singh *et al*., 2016.; Qin *et al*., 2017; Ren *et al*., 2018; Banahene *et al*., 2022). Fluorescence microscopy was used as a semi-quantitative approach, relying on consistent exposure settings and region-of-interest (ROI)-based image analysis to estimate labeling intensity and area. This method is well-suited for detecting spatial and temporal trends in protein modification dynamics during development or disease progression (Hoffmann *et al*. 2022; Hossain *et al*., 2023). For this purpose, 10 µL of the PBS solutions containing L1 and L4 worm stages of N2 and *aex-3*p::tau (V337M) strains were gently mixed and analyzed separately at RT without ambient light throughout the fluorescence microscopy experiment. AD and aging *C. elegans* model images were acquired at 469nm excitation and 525nm emission using an Eclipse TE2000-E inverted fluorescence microscope (Nikon Instruments Inc, Tallahassee, Florida, USA) equipped with a cooled pE-300white Light Source: 77 mm(w) x 186 mm(d) x 162 mm(h), an ORCA Spark CMOS camera (Hamamatsu) and a T-RCP Controller (Nikon). Images were recorded with NIS Elements 3.10 and examined with the Java Image Processing Pipeline (JIPipe) 1.80.0. Image segmentation and tracking were performed using JIPipe v1.80.0 with TrackMate modules integrated via Fiji. All parameter choices—including Gaussian blur (σ = 5), DoG detector thresholds (0.5–1.0), and LAP tracking constraints—were optimized through preliminary testing and guided by validated protocols from the JIPipe user community (Hoffmann *et al*., 2022; Gerst *et al*., 2023). These standardized workflows ensure reproducibility, reduce observer bias, and allow precise tracking of fluorescently labeled structures in complex biological images.

### Image Processing

Image pretreatment was performed by using the image processor Fiji integrated into JIPipe. All experiments were conducted in biological triplicates (three independent experiments on different days for a total of 16 samples). Micrographic replicates for each stage were combined into an 8-bit single file by choosing the File>Import>Image tool in Fiji. Individual images were duplicated and renamed to copy for each trial, consolidated into a single file, and converted to (1900x1200 pixels) to avoid size constraints.

For the background correction, the color was changed to grayscale via the LUT option. Any background was then removed using the Process>Subtract Background command with the rolling ball radius set to the measured pixels and the light background box selected. To make the contrast between the dark background and the worm’s area outstanding, the brightness/contrast was changed (Image>Adjust>Brightness/Contrast). The threshold was then set to the automatically proposed values (Image>Adjust>Threshold), which were between 50 and 250 to obtain representative areas corresponding to the labeled objects (here ROIs). To confirm that the background of the image was changed, we tested different LUTs. Since the green channel seemed to have a very low background, we proceeded to duplicate the image appearance to subtract any left interference by measuring the objects and setting the obtained values in the Analyze>Set Measurements and then Process>Math >Subtract tools box. This new image was saved, and the box labeled “Calculate Threshold for Each Image” was chosen in the “Convert Stack to Binary” box. Pretreated images were saved as a TIFF file for the subsequent analysis (File > Save as > TIFF). (Fig.1A-E)

### Automated Tracking Analysis

Preprocessed files were imported to track labeled spots corresponding to O-GlcNAc by creating a TrackMate project workflow in the visual programming language pipeline JIPipe version 1.80.0 (Research Group Applied Systems Biology, University of Jena, Germany) to open preprocessed .tif files in Fiji. We first followed the step-by-step protocol in the interface for segmentation, which divided the objects through algorithms for displaying and quantifying 2D and 3D images (binary mask detector). Hereby, we selected the 2D mask detector option to identify, and filter labeled spots. This produces an identification of every single object by defining its distinct computing area, shapes and perimeter scores.

For this purpose, we assigned a LUP (Lookup table) to map the spots color as integer pixel values, after Gaussian blurring (Gaussian Blur 2D) parameters were set to sigma X (5) and sigma Y (-1). Following this selection, we modified the estimated blob diameter, thresholding, and median filter in the DoG (Difference of Gaussians) detector box. Subsequently, we chose the median filter and tested the threshold up to 0.5 previously to increase the likelihood that each spot was detected. Finally, we deselected the median filter and decided to set the threshold value to 1, as it avoided including undesirable noise background signals.

Next, we selected a LAP (Linear assignment problem) tracker, set the Gap-closing distance to 15 pixels, a Linking max distance to 15 pixels, a Gap-closing max. frame gap to 2, and filters on tracks with a quality value >3 to detect the most frames (>99%) and significantly reduce noise signals. A “Converting spots to Regions of Interest (ROIs)” node was created and connected to the filter spots box; filters were used to eliminate anomalous ROIs. Total statistics for “All Spots” are performed by JIPipe as well as for adult worms. Statistics for the detected areas were exported after selecting the “Extract Statistics” action box and clicking “Execute”. The spot detection file, automatically produced by TrackMate, was compared with the All-Spots statistics file to make sure that only one spot was marked for each frame and that incorrect duplicates and/or aberrant ROIs were removed. Subsequently, we extracted and plotted estimates of the areas (A), perimeters (P), and circularity (C) of the labeled spots associated with XY trajectories from the analysis tables as a .csv file (Supplementary dataset 1). These allowed us to identify O-GlcNAc labeling to detect biologically relevant changes in protein O-GlcNAcylation, a dynamic post-translational modification regulated by O-GlcNAc transferase (OGT) and O-GlcNAcase (OGA).

### Statistical Analysis

Data are presented as means ± standard deviation (SD) from at least three independent biological replicates, unless otherwise stated. Violin plots, scatter plots, box graphs, and line graphs were created with OriginPro 2025 (Origin Lab Corporation, Northampton, MA, USA). Normality of the data was assessed using the Shapiro-Wilk test. Depending on the distribution of the data, statistical differences were determined using Mann-Whitney-Test pairwise comparisons and Kruskal-Wallis ANOVA with Dunn’s post hoc test. All statistical analyses were conducted in OriginPro 2025. P values of less than .05 were considered statistically significant (*, p < .05; **, p < .01; ***, p < .001). (Supplementary dataset 4)

## Results

### Monitoring Morphological Changes

WT N2 and *aex*-3p::tau (V337M) *C. elegans* strains were cultured and monitored from the young larval stage (L1, day 1) to middle-aged adults (L4, day 11). Fluorescence microscopy images were processed in Fiji, with brightness, threshold, and background adjusted (Fig. 1A-E). O-GlcNAc levels were tracked using labeled regions of interest (ROIs) identified via TrackMate integrated to JIPipe.

Cell morphometric features were quantified based on the detected fluorescence signals. On day 1, shortly after hatching, a substantial number of labeled spots representing newly synthesized O-GlcNAc were observed. we quantified the mean cell area (μm²) across four experimental groups: N2L1 (Group A), N2L4 (Group B), *aex*-3p::tau (V337M) L1 (Group C), and *aex*-3p::tau (V337M) L4 (Group D). The data were visualized using violin plots that display both the distribution of individual data points and the central tendency with error bars representing standard error or standard deviation.

The WT groups, N2L1 and N2L4, showed similar distributions with median labelled areas around 850–900 μm², indicating no statistically significant difference between these two stages (Dunn’s test: N2L1 vs. N2L4, p >.05). The *aex*-3p::tau (V337M) L1 group also showed a comparable distribution (Dunn’s test: N2L4 vs. *aex*-3p::tau (V337M) L1, p > .05, not significant).

In contrast, the *aex*-3p::tau (V337M) L4 group exhibited a striking reduction in mean cell area, with median values centred around 200 μm². This reduction was statistically significant when compared to all other groups: N2 L1 vs. *aex*-3p::tau (V337M) L4; N2 L4 vs. *aex*-3 L4, and *aex*-3 L1 vs. *aex*-3p::tau (V337M) L4: p < .0001. The spread and shape of the violin plots further highlight intra-group variation. Groups B and D, which display broader violin shapes, exhibit higher variability in individual measurements, while Groups A and C are more constrained, indicating less within-group morphological heterogeneity (Fig. 2A-B). The substantial difference in area between *aex*-3:: L1 and *aex*-3p::tau (V337M) L4 (Dunn’s test: 174.83 μm², p <.0001) underscores a developmental stage-specific effect of tau (V337M) mutation, becoming prominent only at the L4 stage (Fig. 2A).

**Figure 2.**
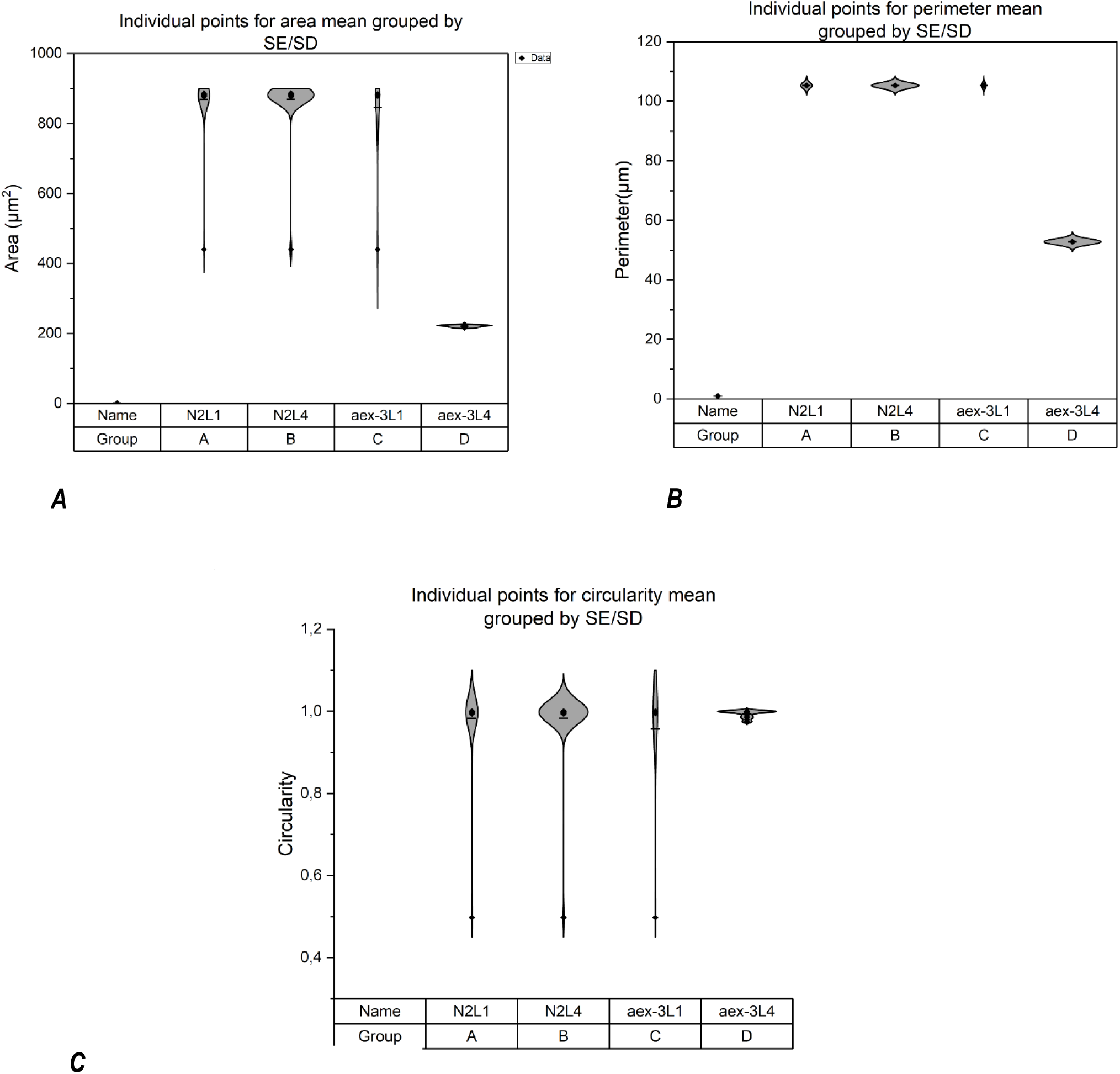
Analysis of morphological parameters and directional angles characterizing the O-GlcNAc labeled spots detected in young and adults N2 and *aex-3*/V337M strains: Violin plots represent the mean ± SE across three experimental biological replicates for a total of n=16 samples. (A) Area, (B) Perimeter and (C) Circularity of the ROIs merged from JIPipe statistics report. Kruskal-Wallis-test of ANOVA (p<0.0001) with Dunńs-test ((p<0.0001) suggest statistically significant differences among spots area sizes (A) and Perimeters (B) for the pairs: N2 L1 - *aex-3*p::tau (V337M) L4, N2 L4 - *aex-3*p::tau (V337M) L4 and *aex-3*p::tau (V337M) L1 and *aex*-3p::tau (V337M) L4 . Statistical analysis for circularity values between 0.5-1.0 (C) suggest a tendency toward a rounded morphological form with insignificant difference of means among strains and life points. (p> 0.05)

Although the number of labeled spots increased in both strains by day 11, the mean perimeter remained relatively consistent for N2 L4 (P = 105.32 µm; please refer to Group B in Fig 2B. In contrast, the *aex-3*p::tau (V337M) L4 exhibited a reduced mean labeled perimeter, representing an approximate twofold decrease compared to day 1 (P ≈ 53 µm). Furthermore, an assumption of normal distribution was rejected for this strain (Shapiro-Wilk test, p < .0001). Therefore, a non-parametric test followed by Dunn’s post hoc test was used for multiple group comparisons. Kruskal-Wallis ANOVA results suggest significant differences in perimeter across middle-aged groups (*p* < .0001), further supported by a post hoc Dunńs HSD test (*p* < .0001).

In the context of TrackMate, circularity quantifies how closely the shape of a detected particle approximates a perfect circle, with a value of 1 indicating a perfect circle (Tinevez *et al*., 2017; Schwendy *et al*., 2019). To assess morphological changes, we analysed circularity scores of O-GlcNAc-labeled regions of interest (ROIs). In the N2L1 group (Group A), circularity values were broadly distributed, ranging from approximately 0.45 to near 1. In contrast, the N2L4 group (Group B) showed a slightly narrower distribution with a higher concentration of values near 1, suggesting that ROIs in this group were generally more circular and morphologically uniform.

The *aex-3*p::tau (V337M) L1 cohort (Group C) displayed a broad circularity spread comparable to N2 L1, but with a slightly higher median value, indicating a partial suppression of the irregular morphologies seen in the control. In contrast, the *aex-3*p::tau (V337M) L4 cohort (Group D) showed the tightest clustering of circularity values around 1, indicating exceptionally uniform, nearly perfect circular shapes. Normality was rejected by the Shapiro– Wilk test (p < .001); however, a Kruskal–Wallis’ test followed by Dunn’s post-hoc comparison found no significant differences among strains or developmental stages (Fig. 2C, p > .05).

### O-GlcNAc Quantification

We measured and analyzed labeled O-GlcNAcylation over a period of 11 days. To ensure accuracy, we examined each area’s value using both raw pixel values and calculated standard errors. Through the reported areas or ROIs at each lifetime point with JIPipe statistical function, pixel-based measurements were analyzed through JIPipés statistical functions, with the percentage of O-GlcNAc computed as:

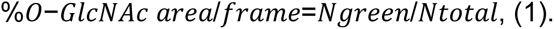

In this equation, Ngreen refers specifically to the number of green pixels identified within the marked ROIs, signifying the presence of O-GlcNAcylation. Ntotal is the total frame area (2,280,000 pixels). Each pixel corresponds to 0.0676 μm^2^, allowing conversion to physical area.

On day 1, early-stage larvae were evaluated. The L1 stage of the N2 strain exhibited a mean of 6.2% O-GlcNAc-labeled area/frame. In contrast, the *aex-3*p::tau (V337M) showed a reduced 2.2% labeled area. This difference was observed despite both strains being at comparable early developmental stages. Repeated analysis of ROIs across young samples indicated a consistent early labeling pattern, with no significant differentiation in spatial distribution at this stage.

By day 11, significant differences emerged between genotypes. In the N2 L4 strain, the mean labelled O-GlcNAc area per frame increased markedly to around 25.4%. In contrast, the transgenic strain expressing human tau (4R1N V337M) under the *aex-3* promoter exhibited only 7.7% of labeled area at the same timepoint, representing a 5.5% reduction relative to its initial day 1 value and an over 3-fold decrease relative to WT N2 at the same developmental stage.

Direct comparison of L4 stage animals on day 11 revealed a strong genotype-dependent effect on O-GlcNAcylation, with N2 exhibiting a 4-fold increase in O-GlcNAc labeled over the transgenic strain. Replicates showed low variability, and trends were consistent across experiments. Kruskal–Wallis’ analysis followed by Dunn’s post hoc test confirmed significant differences in %O-GlcNAcylated area per frame (*p*: .02, please refer to Supplementary Dataset 3).

### Directionality and Migration

To examine the spatial relationship between locomotor directionality and O-GlcNAcylated ROIs, we analysed the 2D trajectories of individual *C. elegans* across developmental stages and genotypes. Scatter plots of movement across the X (lateral) and Y (vertical) axes (Fig. 3A–B) revealed distinct directional behavior that was both genotype- and stage-dependent.

**Figure 3.**
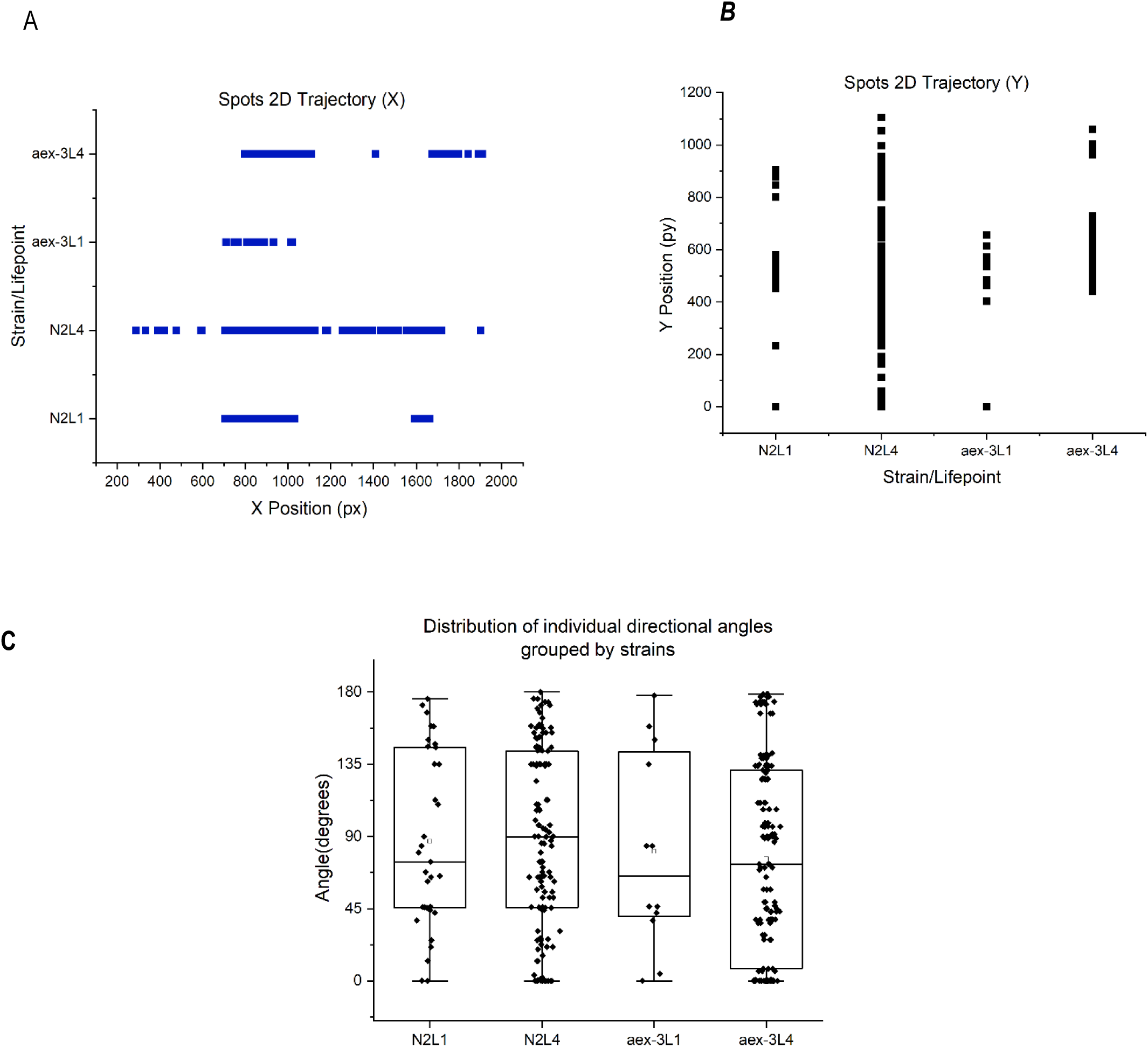
Scatter plots of ROIs level trajectories and distribution of directional angles of labeled O-GlcNAc sites from processed micrographs of an AD and Aging *C. elegans* model. ROIs were selected based on a thresholding between 0,5-1,0 and quantitative data were extracted from JIPipe all spots’ statistics. (A) X-trajectory positions. (B) Y-trajectory positions. The X(lateral) -Y (vertical) positions (in pixels) of detected spots are plotted for four groups: WT N2L1, WT N2L4, *aex-3*p::tau (V337M) L1), and *aex-3*p::tau (V337M) L4). Each blue/black point represents the position of a detected spot over time for individual animals at the corresponding larval stage. (C) Distribution of individual directional angles across strains and developmental stages. Each boxplot represents the spread of angular data, with median lines and overlaid individual data points for every strain. Mann-Whitney Test with Dunńs pot hoc test (p = 0.045) suggest a significant difference in distribution of directional angle values for the comparison pair N2 L4 - *aex-3*p::tau (V337M) L4.

### Lateral Movement and O-GlcNAc Correlation Along the X-Axis

Trajectory analysis along the X-axis (Fig. 3A) showed that WT N2 L1 larvae moved within two distinct horizontal bands (∼730–1000 px and 1500–1600 px), with overlapping paths indicating limited lateral displacement. Correspondingly, O-GlcNAc-labeled ROIs were sparsely distributed, primarily in central regions of the body.

WT L4 larvae exhibited broader lateral displacement, covering nearly the entire imaging field (∼180–1900 px), with frequent directional shifts. This pattern was associated with an increased number and wider distribution of O-GlcNAc-positive ROIs in neuronal and muscular tissues.

In *aex-3*p::tau (V337M) L1 animals, lateral trajectories were confined (∼730–950 px) and discontinuous. O-GlcNAc labeling in these animals was punctate and spatially restricted. The L4 transgenic animals showed modest expansion of lateral range (∼860–1050 px and 1500– 1800 px), but movement remained significantly limited compared to WT L4 animals (Kruskal-Wallis with Dunn’s post hoc test, *p*: .03). O-GlcNAc ROIs in these samples showed limited lateral spread and irregular distribution.

### Vertical Displacement Reflects PTM-Regulated Postural Dynamics

Y-axis movement trajectories (Fig. 3B) in WT N2 L1 larvae were multimodal, with position clusters near 0 px, 450–600 px, and 800–900 px. O-GlcNAc staining was localized to discrete body regions, aligning with positional clustering of ROIs.

In WT L4 larvae, vertical displacement extended continuously from 0 to ∼1100 px, with a more uniform spatial distribution of O-GlcNAc-labeled ROIs throughout the body. *aex-3*p::tau (V337M) L1 animals showed reduced vertical movement, primarily between 0–650 px, with a strong concentration of positions at 0 px. O-GlcNAc labeling in these animals was restricted to perinuclear zones. The L4 transgenic group exhibited moderate vertical displacement (∼420–1050 px); however, tracks remained fragmented, and ROI localization lacked the continuity observed in WT L4 samples (p < .0001, Kruskal-Wallis with Dunn’s test).

### Directional angles

To further clarify these observations, we classified the directional angle values of the observed trajectories as anterograde (< 90°) or retrograde (> 90°), with angles ≤ 1° treated as an additional near-0° “non-directional or stationary points”. The distribution of these categories was then compared between WT N2 and *aex-3*p::tau (V337M) strains at two developmental stages (young L1 and middle-aged L4). A summary of the percentages is provided in Table S1.

In WT N2 L1 larvae, O-GlcNAc-modified ROIs showed a clear forward bias: 49.6 % of trajectories were anterograde, versus 38.5 % retrograde. In striking contrast, for *aex-3*p::tau V337M L1 larvae were predominantly retrograde-oriented; only 3.5 % of movements fell below 90°, indicating a severe loss of forward-directed motion and suggesting early mis-routing or exploratory instability.

Directional complexity increased in both strains. O-GlcNAcylated ROIs in N2 L4 animals retained a modest anterograde predominance (55.4 % anterograde, 42.4 % retrograde) with a small subset (13.0 %) of near-0° paths, consistent with the balanced but purposeful locomotion typical of middle-aged larval exploration. In *aex-3*p::tau V337M animals, the anterograde fraction increased to 46.7 % but stayed below the WT cohort, retrograde motion persisted at 44.8 %, and the proportion of near-0° trajectories nearly doubled to 19.8 %. These abrupt directional switches were most pronounced in the head and anterior body segments. Pairwise comparisons of Mann-Whitney test suggest a significant difference in directional angle distributions between WT N2 and *aex-3*p::tau (V337M) on day 11 (p: .04; see Fig. 3C and Supplementary dataset 2).

## Discussion

In this study, we investigate phenotypic changes associated with O-GlcNAcylation in a *C. elegans* model of Alzheimer’s disease and aging. By using a fluorescence chemoenzymatic approach combined with a quantitative bioimage analysis pipeline, we examined O-glycosylation dynamics and organismal performance across developmental stages. Our results revealed three key effects: (i) a pronounced reduction in O-GlcNAc–positive cell area and perimeter that appears specifically at the L4 larval stage in the transgenic strain expressing human tau (4R1N V337M) under the *aex-3* promoter, (ii) an absence of the age-related increase in O-GlcNAcylation seen in WT animals, and (iii) a shift in locomotor patterns from the forward-biased repertoire typical of WT N2 worms to a fragmented, retrograde-dominated movement in transgenic lines. Despite a modest sample size dataset, the convergence of findings across independent measurements supports the interpretation that these effects represent meaningful phenotypic outcomes rather than random variation.

In terms of morphometric changes, WT N2 strain of *C. elegans* maintained relatively stable regions of interest (ROIs) from the L1 to L4 stages, whereas tau(V337M) transgenic animals exhibited an approximately fourfold reduction in ROI area by day 11. This morphological shrinkage during the late larval-to-adult transition mirrors cytoskeletal destabilization observed in other V337M tauopathy models, where impaired microtubule binding leads to axonal degeneration (Sohn *et al*., 2019). Since ROIs were defined using chemoenzymatic labeling to detect O-GlcNAc-modified structures, the observed reduction likely reflects a loss of glycosylated cellular compartments. Whether this decrease represents genuine cell atrophy or reduced accessibility of O-GlcNAc domains requires confirmation through ultrastructural analysis; however, either interpretation aligns with prior reports that O-GlcNAc modification contributes to cytoskeletal integrity (McColgan *et al*., 2020) and that depletion of neuronal OGT accelerates dendritic retraction in mouse models (Dong *et al*., 2023).

Interestingly, no significant differences in circular morphology were observed between strains or life stages, with a median circularity remaining above 0.99. This suggests that O-GlcNAcylation may not directly correlate with morphological changes in our model. Instead, we propose that O-GlcNAc-modified regions predominantly retained compact, non-elongated shapes throughout development, consistent with prior findings that link O-GlcNAc modification to nuclear and cytoplasmic protein scaffolds (Eustice *et al*., 2017; Chatham *et al*., 2021). This contrasts with the findings of Pinho *et al*., who linked O-GlcNAc loss to morphological disruption in post-mortem AD brain tissue. However, their data reflect a terminal disease state, whereas our *in vivo* model may better represent dynamic and potentially reversible changes during aging. Although O-GlcNAcylation has been implicated in morphological regeneration in *C. elegans* (Taub *et al*., 2018), its role in maintaining cell shape across the lifespan remains to be elucidated.

In WT N2 worms, O-GlcNAc labeling increased approximately fourfold from day 1 to day 11, consistent with studies showing that O-GlcNAcylation rises during aging to buffer proteotoxic stress (Mueller *et al*., 2021). In contrast, the transgenic strain failed to show this adaptive increase and showed declining O-GlcNAc levels with age. This impaired response may stem from competition between mutant tau and O-GlcNAc transferase (OGT) for sugar donors or from steric interference with OGT access, in line with evidence of reciprocal regulation between tau phosphorylation and O-GlcNAcylation (Tramutola *et al*., 2018). Given that all animals were kept in nutrient-rich conditions, we hypothesize that metabolic, trafficking or enzymatic alterations due to aging or AD mutations affect O-GlcNAc patterns. This observation aligns with the results reported by Pinho *et al*., 2019, who showed reduced global O-GlcNAcylation in an *in vitro* AD model.

Behaviorally, trajectory analysis revealed that V337M-expressing worms exhibited a marked bias toward retrograde movement from the L1 stage onward, with further exaggeration at L4. This retrograde dominance echoes the uncoordinated phenotype of established *C. elegans* tauopathy models and suggests disrupted neuromuscular coordination. Importantly, the degree of locomotor disorganization correlated with the distribution of O-GlcNAc–positive regions: worms with broader movement ranges had abundant labeling, while those with more restricted movement showed sparse or punctate labeling. Given the known roles of OGT and OGA in axonal regeneration and dauer formation, a causal relationship between sugar signaling and motor behavior is plausible (Wu *et al*., 2024; Lee *et al*., 2021). This may point to a bidirectional relationship, where altered motility impacts O-GlcNAc cycling, which in turn feeds back into cellular transport or signaling pathways. However, establishing causality will require targeted manipulations of O-GlcNAc transferase or hydrolase activity in specific neuronal populations.

Our findings align with prior studies linking impaired intracellular trafficking to neurodegeneration. Notably, Sleigh *et al*. (2017) and Mandal *et al*. (2021) reported that aging and disease states can selectively disrupt anterograde transport, reducing the delivery of essential organelles and synaptic proteins. In the context of O-GlcNAc signaling, reduced anterograde dynamics may indicate compromised support for synaptic maintenance. Conversely, a relative increase in retrograde transport could reflect a shift toward the clearance of dysfunctional cellular components, potentially as a compensatory mechanism in later developmental stages. This shift in trafficking polarity highlights the possibility that disrupted O-GlcNAc cycling contributes to a broader failure in neuronal homeostasis, particularly in the *aex-3*p::tau (V337M) background.

The presence of non-directional alignment (near 0°) may reflect underlying processes related to cellular maintenance or regeneration. While more frequent in the *aex-3* L4 transgenic strain compared to WT N2, such alignment patterns were also detected in early-stage starved *aex-3* L1 animals. Previous studies have associated low-angle alignment with cytoskeletal remodeling and regenerative responses in muscle and neural tissues (Suarez *et al*., 2019; Drewry *et al*., 2022). Although the mechanistic role of O-GlcNAcylation in this context remains to be clarified, reports linking modified O-GlcNAc levels to enhanced regeneration in *ogt-1* mutants (Taub *et al*., 2018; Yadav *et al*., 2024) suggest it may influence the cellular capacity to adapt under stress. These observations support the idea that altered O-GlcNAc signaling could contribute to dynamic changes in cytoskeletal behavior during development or repair, particularly in genetically susceptible backgrounds.

Our findings add to previous work showing that O-GlcNAc depletion exacerbates tau pathology in mammals by demonstrating a similar interaction *in vivo* in nematodes. Notably, this study is among the first to quantitatively link directional locomotion with O-GlcNAc distribution using JIPipe-based analysis, providing insights into the dynamics of this PTM in neurodegenerative contexts. The obtained labeling intensities reflect underlying biological differences in enzyme activity, metabolic flux through the hexosamine biosynthetic pathway, and cellular stress responses, rather than simple variations in fluorescence signal. Thus, differences in O-GlcNAc labeling are interpreted as indicative of altered cellular physiology, consistent with previous studies linking O-GlcNAc dynamics to aging and neurodegenerative processes (Ortega-Prieto *et al*., 2023; Costa *et al*., 2023).

We also recognize several limitations. First, the sample size restricts statistical power, necessitating the use of non-parametric analyses, which may overlook subtler trends. Second, only one transgenic line was analysed, raising the possibility of insertion-site effects. A validation of findings with additional independent lines of *C. elegans* expressing human wild-type and mutant tau will be essential for generalization.

To build on these preliminary findings, we propose several next steps. Increasing the sample size to at least 48 worms would enable more robust mixed-effects modelling. Including later developmental stages (beyond day 15) could help determine whether motor phenotypes progress to paralysis. Proteomic analysis to study hypoglycosylation and O-GlcNAc sites on biomarkers that could link O-GlcNAc dynamics to neural activity patterns that underlie directional choices in AD. These proposed directions are directly grounded in the observed data and avoid vague generalities.

In summary, our results suggest that impaired O-GlcNAc upregulation during development increases neuronal vulnerability to mutant tau, leading to structural degeneration and altered motor behavior. While exploratory, this study illustrates the power of bioimaging pipelines to examine this post-translational modification and offers a quantitative framework for behavioral phenotyping in *C. elegans* neurodegeneration models.

## Acknowledgments

We thank Dr. Edmund Kostewicz for helpful discussions and feedback on the manuscript; Dr. Maximilian Pfisterer and Prof. Dr. Lienhard Schmitz from the Biochemistry Institute of the University of Giessen for the guidance on the fluorescence microscopy method. *aex-3*p::4R1N-Tau(V337M) strain was kindly provided by Dr. Brian C. Kraemer from the UW Division of Gerontology & Geriatric Medicine, Geriatric Research, Education and Clinical Center (GRECC), Department of Medicine, and Division of Neurogenetics, Department of Neurology, University of Washington, Seattle, WA, USA.

## Data and resource availability

JIPipe-processed spot tracking reports and Origin 2025 statistics are provided as Supplementary Data Files 1-4. Additional data available upon request.

