## Supplementary material for "Tracking of *in vivo* O-GlcNAcylation in an Alzheimer’s Disease and Aging *C. elegans* Model": Pair-B-C-comparison.pdf

### Mann-Whitney Test (29/06/2025 15:59:35)

#### Notes

| | |
|---|---|
| X-Function | Mann-Whitney Test |
| User Name | fgaro |
| Time | 29/06/2025 15:59:35 |
| Data Filter | No |

#### Input Data

| | Data | Range |
|---|---|---|
| 1st Data Range | [Celegans]Angles!B" | [1*:136*] |
| 2nd Data Range | [Celegans]Angles!C" | [1*:13*] |

#### Descriptive Statistics

| | N | Min | Q1 | Median | Q3 | Max |
|---|---|---|---|---|---|---|
| B | 135 | 0 | 45,564 | 89,738 | 143,28 | 180 |
| C | 12 | 0 | 38,87075 | 65,2085 | 146,421 | 177,849 |

#### Ranks

| | N | Mean Rank | Sum Rank |
|---|---|---|---|
| B | 135 | 74,47778 | 10054,5 |
| C | 12 | 68,625 | 823,5 |

#### Test Statistics

| U | Z | Asymp. Prob>\|U\| |
|---|---|---|
| 874,5 | 0,45316 | 0,65044 |

Null Hypothesis: Median1 = Median2

Alternative Hypothesis: Median1 <> Median2

**At the 0.05 level, the two distributions are NOT significantly different.**
