## Supplementary material for "Tracking of *in vivo* O-GlcNAcylation in an Alzheimer’s Disease and Aging *C. elegans* Model": Pair_A-B-comparison.pdf

### Mann-Whitney Test (29/06/2025 15:28:20)

#### Notes

| | |
|---|---|
| X-Function | Mann-Whitney Test |
| User Name | fgaro |
| Time | 29/06/2025 15:28:20 |
| Data Filter | No |

#### Input Data

| | Data | Range |
|---|---|---|
| 1st Data Range | [Celegans]Angles!A" | [1*:34*] |
| 2nd Data Range | [Celegans]Angles!B" | [1*:136*] |

#### Descriptive Statistics

| | N | Min | Q1 | Median | Q3 | Max |
|---|---|---|---|---|---|---|
| A | 33 | 0 | 44,806 | 74,19 | 145,7 | 175,831 |
| B | 135 | 0 | 45,564 | 89,738 | 143,28 | 180 |

#### Ranks

| | N | Mean Rank | Sum Rank |
|---|---|---|---|
| A | 33 | 83,45455 | 2754 |
| B | 135 | 84,75556 | 11442 |

#### Test Statistics

| U | Z | Asymp. Prob>\|U\| |
|---|---|---|
| 2193 | -0,13584 | 0,89195 |

Null Hypothesis: Median1 = Median2

Alternative Hypothesis: Median1 <> Median2

**At the 0.05 level, the two distributions are NOT significantly different.**
