## Supplementary material for "Tracking of *in vivo* O-GlcNAcylation in an Alzheimer’s Disease and Aging *C. elegans* Model": Pair_A-C-comparison.pdf

### Mann-Whitney Test (29/06/2025 15:29:59)

#### Notes

| | |
|---|---|
| X-Function | Mann-Whitney Test |
| User Name | fgaro |
| Time | 29/06/2025 15:29:59 |
| Data Filter | No |

#### Input Data

| | **Data** | **Range** |
|---|---|---|
| 1st Data Range | [Celegans]Angles!A" | [1*:34*] |
| 2nd Data Range | [Celegans]Angles!C" | [1*:13*] |

#### Descriptive Statistics

| | **N** | **Min** | **Q1** | **Median** | **Q3** | **Max** |
|---|---|---|---|---|---|---|
| A | 33 | 0 | 44,806 | 74,19 | 145,7 | 175,831 |
| C | 12 | 0 | 38,87075 | 65,2085 | 146,421 | 177,849 |

#### Ranks

| | **N** | **Mean Rank** | **Sum Rank** |
|---|---|---|---|
| A | 33 | 23,33333 | 770 |
| C | 12 | 22,08333 | 265 |

#### Test Statistics

| **U** | **Z** | **Asymp. Prob>\|U\|** |
|---|---|---|
| 209 | 0,26967 | 0,78741 |

Null Hypothesis: Median1 = Median2

Alternative Hypothesis: Median1 <> Median2

**At the 0.05 level, the two distributions are NOT significantly different.**
