## Supplementary material for "Tracking of *in vivo* O-GlcNAcylation in an Alzheimer’s Disease and Aging *C. elegans* Model": Pair_A-D-comparison.pdf

### Mann-Whitney Test (29/06/2025 15:31:10)

#### Notes

| | |
|---|---|
| X-Function | Mann-Whitney Test |
| User Name | fgaro |
| Time | 29/06/2025 15:31:10 |
| Data Filter | No |

#### Input Data

| | Data | Range |
|---|---|---|
| 1st Data Range | [Celegans]Angles!A" | [1*:34*] |
| 2nd Data Range | [Celegans]Angles!D" | [1*:163*] |

#### Descriptive Statistics

| | N | Min | Q1 | Median | Q3 | Max |
|---|---|---|---|---|---|---|
| A | 33 | 0 | 44,806 | 74,19 | 145,7 | 175,831 |
| D | 162 | 0 | 7,58775 | 72,73 | 131,347 | 178,7 |

#### Ranks

| | N | Mean Rank | Sum Rank |
|---|---|---|---|
| A | 33 | 108,28788 | 3573,5 |
| D | 162 | 95,90432 | 15536,5 |

#### Test Statistics

| U | Z | Asymp. Prob>\|U\| |
|---|---|---|
| 3012,5 | 1,15063 | 0,24989 |

Null Hypothesis: Median1 = Median2

Alternative Hypothesis: Median1 <> Median2

**At the 0.05 level, the two distributions are NOT significantly different.**
