## Supplementary material for "Tracking of *in vivo* O-GlcNAcylation in an Alzheimer’s Disease and Aging *C. elegans* Model": Pair_B-D-comparison.pdf

### Mann-Whitney Test (29/06/2025 15:16:34)

#### Notes

| | |
|---|---|
| X-Function | Mann-Whitney Test |
| User Name | fgaro |
| Time | 29/06/2025 15:16:34 |
| Data Filter | No |

#### Input Data

| | Data | Range |
|---|---|---|
| 1st Data Range | [Celegans]Angles!B" | [1*:136*] |
| 2nd Data Range | [Celegans]Angles!D" | [1*:163*] |

#### Descriptive Statistics

| | N | Min | Q1 | Median | Q3 | Max |
|---|---|---|---|---|---|---|
| B | 135 | 0 | 45,564 | 89,738 | 143,28 | 180 |
| D | 162 | 0 | 7,58775 | 72,73 | 131,347 | 178,7 |

#### Ranks

| | N | Mean Rank | Sum Rank |
|---|---|---|---|
| B | 135 | 159,8963 | 21586 |
| D | 162 | 139,91975 | 22667 |

#### Test Statistics

| U | Z | Asymp. Prob>\|U\| |
|---|---|---|
| 12406 | 1,99922 | 0,04558 |

Null Hypothesis: Median1 = Median2

Alternative Hypothesis: Median1 <> Median2

**At the 0.05 level, the two distributions are significantly different.**
