## Supplementary material for "Tracking of *in vivo* O-GlcNAcylation in an Alzheimer’s Disease and Aging *C. elegans* Model": Pair_C-D-comparison.pdf

### Mann-Whitney Test (29/06/2025 16:00:16)

#### Notes

| | |
|---|---|
| X-Function | Mann-Whitney Test |
| User Name | fgaro |
| Time | 29/06/2025 16:00:16 |
| Data Filter | No |

#### Input Data

| | Data | Range |
|---|---|---|
| 1st Data Range | [Celegans]Angles!C" | [1*:13*] |
| 2nd Data Range | [Celegans]Angles!D" | [1*:163*] |

#### Descriptive Statistics

| | N | Min | Q1 | Median | Q3 | Max |
|---|---|---|---|---|---|---|
| C | 12 | 0 | 38,87075 | 65,2085 | 146,421 | 177,849 |
| D | 162 | 0 | 7,58775 | 72,73 | 131,347 | 178,7 |

#### Ranks

| | N | Mean Rank | Sum Rank |
|---|---|---|---|
| C | 12 | 92,625 | 1111,5 |
| D | 162 | 87,12037 | 14113,5 |

#### Test Statistics

<table>
<tr><th>U</th><th>Z</th><th>Asymp. Prob>|U|</th></tr>
<tr><td>1033,5</td><td>0,36367</td><td>0,7161</td></tr>
</table>

Null Hypothesis: Median1 = Median2

Alternative Hypothesis: Median1 <> Median2

**At the 0.05 level, the two distributions are NOT significantly different.**
