## Supplementary material for "Tracking of *in vivo* O-GlcNAcylation in an Alzheimer’s Disease and Aging *C. elegans* Model": posthoc_2groups_B_D.pdf

### Kruskal-Wallis ANOVA (09/07/2025 14:15:09)

#### Notes

| | |
|---|---|
| X-Function | Kruskal-Wallis ANOVA |
| User Name | fgaro |
| Time | 09/07/2025 14:15:09 |
| Data Filter | No |

#### Input Data

| | Data | Range |
|---|---|---|
| Group Range | [Celegans]Angles!A"N2L4" | [1*:298*] |
| Data Range | [Celegans]Angles!B | [1*:298*] |

#### Descriptive Statistics

| | | N | Min | Q1 | Median | Q3 | Max |
|---|---|---|---|---|---|---|---|
| B | N2L4 | 135 | 0 | 45,564 | 89,738 | 143,28 | 180 |
| | aex-3L4 | 162 | 0 | 7,58775 | 72,73 | 131,347 | 178,7 |

#### Ranks

| | | N | Mean Rank | Sum Rank |
|---|---|---|---|---|
| B | N2L4 | 135 | 159,8963 | 21586 |
| | aex-3L4 | 162 | 139,91975 | 22667 |

#### Test Statistics

| | Chi-Square | DF | Prob>Chi-Square |
|---|---|---|---|
| B | 3,9996 | 1 | 0,04551 |

Null Hypothesis:The samples come from the same population.
Alternative Hypothesis:The samples come from different populations.

**B: At the 0.05 level, the populations are significantly different.**

#### Dunn's Test

| | | | Mean Rank Diff | Z | Prob | Sig |
|---|---|---|---|---|---|---|
| B | N2L4 | aex-3L4 | 19,97654 | 1,9999 | 0,04551 | 1 |

Sig equals 1 indicates that the difference of the means is significant at the 0,05 level.
Sig equals 0 indicates that the difference of the means is NOT significant at the 0,05 level.
