## Supplementary material for "Tracking of *in vivo* O-GlcNAcylation in an Alzheimer’s Disease and Aging *C. elegans* Model": posthoc_OGlcNAcquantification.pdf

### Kruskal-Wallis ANOVA (27/06/2025 11:09:31)

#### Notes

| | |
|---|---|
| X-Function | Kruskal-Wallis ANOVA |
| User Name | fgaro |
| Time | 27/06/2025 11:09:31 |
| Data Filter | No |

#### Input Data

| | Data | Range |
|---|---|---|
| Group Range | [Celegans]Sheet3!A | [1*:365*] |
| Data Range | [Celegans]Sheet3!C | [1*:366*] |

#### Descriptive Statistics

| | | N | Min | Q1 | Median | Q3 | Max |
|---|---|---|---|---|---|---|---|
| C | N2L1 | 3 | 2,9 | 2,9 | 4,3 | 11,4 | 11,4 |
| | N2L4 | 3 | 18,3 | 18,3 | 24,6 | 33,2 | 33,2 |
| | aex-3L1 | 3 | 1,43 | 1,43 | 1,7 | 3,4 | 3,4 |
| | aex-3L4 | 3 | 6,9 | 6,9 | 7,9 | 8,5 | 8,5 |

#### Ranks

| | | N | Mean Rank | Sum Rank |
|---|---|---|---|---|
| C | N2L1 | 3 | 5,66667 | 17 |
| | N2L4 | 3 | 11 | 33 |
| | aex-3L1 | 3 | 2,33333 | 7 |
| | aex-3L4 | 3 | 7 | 21 |

#### Test Statistics

| | Chi-Square | DF | Prob>Chi-Square |
|---|---|---|---|
| C | 8,89744 | 3 | 0,03069 |

Null Hypothesis:The samples come from the same population.
Alternative Hypothesis:The samples come from different populations.

**C: At the 0.05 level, the populations are significantly different.**

#### Dunn's Test

| | | | Mean Rank Diff | Z | Prob | Sig |
|---|---|---|---|---|---|---|
| C | N2L1 | N2L4 | -5,33333 | -1,81164 | 0,42025 | 0 |
| | N2L1 | aex-3L1 | 3,33333 | 1,13228 | 1 | 0 |
| | N2L1 | aex-3L4 | -1,33333 | -0,45291 | 1 | 0 |
| | N2L4 | aex-3L1 | 8,66667 | 2,94392 | 0,01945 | 1 |
| | N2L4 | aex-3L4 | 4 | 1,35873 | 1 | 0 |
| | aex-3L1 | aex-3L4 | -4,66667 | -1,58519 | 0,67754 | 0 |

Sig equals 1 indicates that the difference of the means is significant at the 0,05 level.
Sig equals 0 indicates that the difference of the means is NOT significant at the 0,05 level.

#### Paired Comparison Plot

C

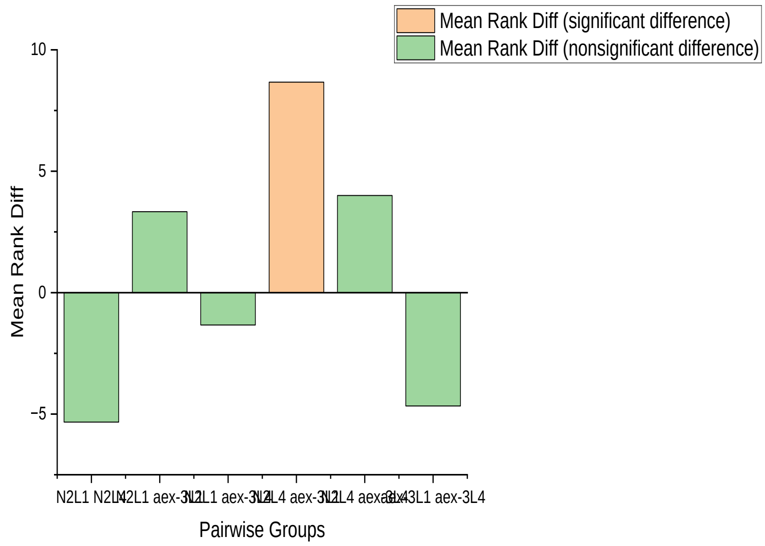
