## Supplementary material for "Tracking of *in vivo* O-GlcNAcylation in an Alzheimer’s Disease and Aging *C. elegans* Model": Norm_test(x-trajectories).pdf

### Normality Test (25/06/2025 13:35:53)

#### Descriptive Statistics

| | | N Analysis | N Missing | Mean | Standard Deviation | SE of Mean |
|---|---|---|---|---|---|---|
| E | Name | 0 | 1 | -- | -- | -- |
| | N2L1 | 33 | 0 | 948,12121 | 275,78997 | 48,00887 |
| | N2L4 | 135 | 0 | 1036,12593 | 355,14482 | 30,566 |
| | aex-3L1 | 12 | 0 | 842,08333 | 104,81886 | 30,2586 |
| | aex-3L4 | 162 | 0 | 1055,82716 | 220,2109 | 17,3014 |

#### NormalityTest

##### Shapiro-Wilk

| | | DF | Statistic | p-value | Decision at level(5%) |
|---|---|---|---|---|---|
| E | Name | -- | -- | -- | a* |
| | N2L1 | 33 | 0,71679 | <0.0001 | Reject normality |
| | N2L4 | 135 | 0,94985 | <0.0001 | Reject normality |
| | aex-3L1 | 12 | 0,91368 | 0,23776 | Can't reject normality |
| | aex-3L4 | 162 | 0,68219 | <0.0001 | Reject normality |

**a*: Too few data points, data points number should not less than 3 for Shapiro-Wilk.**
**E(N2L1): At the 0.05 level, the data was not significantly drawn from a normally distributed population.**
**E(N2L4): At the 0.05 level, the data was not significantly drawn from a normally distributed population.**
**E(aex-3L1): At the 0.05 level, the data was significantly drawn from a normally distributed population.**
**E(aex-3L4): At the 0.05 level, the data was not significantly drawn from a normally distributed population.**
