## Supplementary material for "Tracking of *in vivo* O-GlcNAcylation in an Alzheimer’s Disease and Aging *C. elegans* Model": Norm_Test(y-trajectories).pdf

### Normality Test (25/06/2025 13:37:36)

#### Notes

| | |
|---|---|
| Description | Perform Normality Test |
| User Name | fgaro |
| Operation Time | 25/06/2025 13:37:36 |
| Report Status | New Analysis Report |
| Data Filter | No |

#### Input Data

| | Data | Range |
|---|---|---|
| Data | [Celegans]Sheet1!F | [1*:343*] |
| Group | [Celegans]Sheet1!B | [1*:343*] |

There exist missing values in the input data.

#### Descriptive Statistics

| | | N Analysis | N Missing | Mean | Standard Deviation | SE of Mean |
|---|---|---|---|---|---|---|
| F | Name | 0 | 1 | -- | -- | -- |
| | N2L1 | 33 | 0 | 532,72727 | 163,2043 | 28,41022 |
| | N2L4 | 135 | 0 | 510,82222 | 243,81088 | 20,9839 |
| | aex-3L1 | 12 | 0 | 487,33333 | 168,36612 | 48,60311 |
| | aex-3L4 | 162 | 0 | 654,24074 | 119,64699 | 9,40036 |

#### NormalityTest

##### Shapiro-Wilk

| | | DF | Statistic | p-value | Decision at level(5%) |
|---|---|---|---|---|---|
| F | Name | -- | -- | -- | a* |
| | N2L1 | 33 | 0,79634 | <0.0001 | Reject normality |
| | N2L4 | 135 | 0,97607 | 0,01762 | Reject normality |
| | aex-3L1 | 12 | 0,72613 | 0,00152 | Reject normality |
| | aex-3L4 | 162 | 0,85817 | <0.0001 | Reject normality |

**a*: Too few data points, data points number should not less than 3 for Shapiro-Wilk.**
**F(N2L1): At the 0.05 level, the data was not significantly drawn from a normally distributed population.**
**F(N2L4): At the 0.05 level, the data was not significantly drawn from a normally distributed population.**
**F(aex-3L1): At the 0.05 level, the data was not significantly drawn from a normally distributed population.**
**F(aex-3L4): At the 0.05 level, the data was not significantly drawn from a normally distributed population.**
