## Supplementary material for "Tracking of *in vivo* O-GlcNAcylation in an Alzheimer’s Disease and Aging *C. elegans* Model": Norm_Test_Area.pdf

### Normality Test (13/06/2025 15:52:22)

#### Notes

| | |
|---|---|
| Description | Perform Normality Test |
| User Name | fgaro |
| Operation Time | 13/06/2025 15:52:22 |
| Report Status | New Analysis Report |
| Data Filter | No |

#### Input Data

| | Data | Range |
|---|---|---|
| Data | [Celegans]Sheet1!C | [6:6] |
| Group | [Celegans]Sheet1!A | [1*:343*] |
| | [Celegans]Sheet1!B | [1*:343*] |

There exist missing values in the input data.

#### Descriptive Statistics

| | | | N Analysis | N Missing | Mean | Standard Deviation | SE of Mean |
|---|---|---|---|---|---|---|---|
| C | Group | Name | 0 | 1 | -- | -- | -- |
| | A | N2L1 | 33 | 0 | 868,36364 | 76,94837 | 13,39499 |
| | B | N2L4 | 135 | 0 | 869,11852 | 75,31229 | 6,48185 |
| | C | aex-3L1 | 12 | 0 | 845,75 | 127,79751 | 36,89196 |
| | D | aex-3L4 | 162 | 0 | 221,1358 | 2,84923 | 0,22386 |

#### NormalityTest

##### Shapiro-Wilk

| | | | DF | Statistic | p-value | Decision at level(5%) |
|---|---|---|---|---|---|---|
| C | Group | Name | -- | -- | -- | a* |
| | A | N2L1 | 33 | 0,19497 | <0.0001 | Reject normality |
| | B | N2L4 | 135 | 0,18539 | <0.0001 | Reject normality |
| | C | aex-3L1 | 12 | 0,34167 | <0.0001 | Reject normality |
| | D | aex-3L4 | 162 | 0,93562 | <0.0001 | Reject normality |

**a*: Too few data points, data points number should not less than 3 for Shapiro-Wilk.**
**C(A,N2L1): At the 0.05 level, the data was not significantly drawn from a normally distributed population.**
**C(B,N2L4): At the 0.05 level, the data was not significantly drawn from a normally distributed population.**
**C(C,aex-3L1): At the 0.05 level, the data was not significantly drawn from a normally distributed population.**
**C(D,aex-3L4): At the 0.05 level, the data was not significantly drawn from a normally distributed population.**
