## Supplementary material for "Tracking of *in vivo* O-GlcNAcylation in an Alzheimer’s Disease and Aging *C. elegans* Model": Norm_Test_Perimeter.pdf

### Normality Test (19/06/2025 16:27:37)

#### Notes

| | |
|---|---|
| Description | Perform Normality Test |
| User Name | fgaro |
| Operation Time | 19/06/2025 16:27:37 |
| Report Status | New Analysis Report |
| Data Filter | No |

#### Input Data

| | Data | Range |
|---|---|---|
| Data | [Celegans]Sheet1!D | [1*:343*] |
| Group | [Celegans]Sheet1!A | [1*:343*] |
| | [Celegans]Sheet1!B | [1*:343*] |

There exist missing values in the input data.

#### Descriptive Statistics

| | | | N Analysis | N Missing | Mean | Standard Deviation | SE of Mean |
|---|---|---|---|---|---|---|---|
| D | Group | Name | 0 | 1 | -- | -- | -- |
| | A | N2L1 | 33 | 0 | 105,32 | 0 | 0 |
| | B | N2L4 | 135 | 0 | 105,32 | 0 | 0 |
| | C | aex-3L1 | 12 | 0 | 105,32 | 0 | 0 |
| | D | aex-3L4 | 162 | 0 | 52,7503 | 4,58039E-4 | 3,5987E-5 |

#### NormalityTest

##### Shapiro-Wilk

| | | | DF | Statistic | p-value | Decision at level(5%) |
|---|---|---|---|---|---|---|
| D | Group | Name | -- | -- | -- | a* |
| | A | N2L1 | 33 | -- | -- | c* |
| | B | N2L4 | 135 | -- | -- | c* |
| | C | aex-3L1 | 12 | -- | -- | c* |
| | D | aex-3L4 | 162 | 0,57302 | <0.0001 | Reject normality |

**a*: Too few data points, data points number should not less than 3 for Shapiro-Wilk.**
**c*: There is not enough information to draw conclusion**
**D(D,aex-3L4): At the 0.05 level, the data was not significantly drawn from a normally distributed population.**
