## Supplementary material for "Tracking of *in vivo* O-GlcNAcylation in an Alzheimer’s Disease and Aging *C. elegans* Model": post_hoc_Anallysis(Cricularity).pdf

### Kruskal-Wallis ANOVA (25/06/2025 09:58:41)

#### Notes

| | |
|---|---|
| X-Function | Kruskal-Wallis ANOVA |
| User Name | fgaro |
| Time | 25/06/2025 09:58:41 |
| Data Filter | No |

#### Input Data

| | Data | Range |
|---|---|---|
| Group Range | [Celegans]Sheet1!B | [1*:343*] |
| Data Range | [Celegans]Sheet1!H | [1*:343*] |

#### Descriptive Statistics

| | | N | Min | Q1 | Median | Q3 | Max |
|---|---|---|---|---|---|---|---|
| H | N2L1 | 33 | 0,498 | 0,996 | 0,999 | 1 | 1 |
| | N2L4 | 135 | 0,498 | 0,996 | 0,999 | 1 | 1 |
| | aex-3L1 | 12 | 0,498 | 0,99675 | 1 | 1 | 1 |
| | aex-3L4 | 162 | 0,975 | 0,989 | 0,99902 | 1 | 1 |

#### Ranks

| | | N | Mean Rank | Sum Rank |
|---|---|---|---|---|
| H | N2L1 | 33 | 173,13636 | 5713,5 |
| | N2L4 | 135 | 179,32963 | 24209,5 |
| | aex-3L1 | 12 | 194,33333 | 2332 |
| | aex-3L4 | 162 | 162,95062 | 26398 |

#### Test Statistics

| | Chi-Square | DF | Prob>Chi-Square |
|---|---|---|---|
| H | 3,05647 | 3 | 0,383 |

Null Hypothesis:The samples come from the same population.
Alternative Hypothesis:The samples come from different populations.
**H: At the 0.0001 level, the populations are NOT significantly different.**

#### Dunn's Test

| | | | Mean Rank Diff | Z | Prob | Sig |
|---|---|---|---|---|---|---|
| H | N2L1 | N2L4 | -6,19327 | -0,3229 | 1 | 0 |
| | N2L1 | aex-3L1 | -21,19697 | -0,63664 | 1 | 0 |
| | N2L1 | aex-3L4 | 10,18575 | 0,53997 | 1 | 0 |
| | N2L4 | aex-3L1 | -15,0037 | -0,50429 | 1 | 0 |
| | N2L4 | aex-3L4 | 16,37901 | 1,42303 | 0,92837 | 0 |
| | aex-3L1 | aex-3L4 | 31,38272 | 1,06205 | 1 | 0 |

Sig equals 1 indicates that the difference of the means is significant at the 0,05 level.
Sig equals 0 indicates that the difference of the means is NOT significant at the 0,05 level.

#### Paired Comparison Plot

H

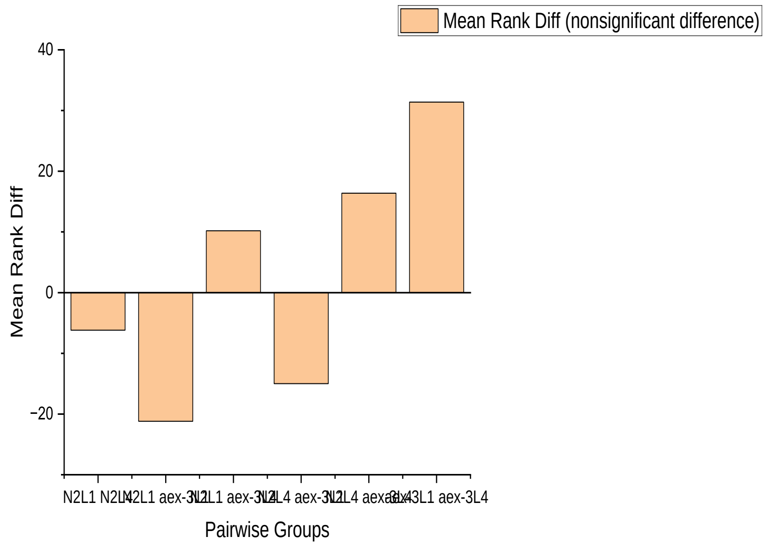
