## Supplementary material for "Tracking of *in vivo* O-GlcNAcylation in an Alzheimer’s Disease and Aging *C. elegans* Model": Post_hoc_Analysis(Area).pdf

### Kruskal-Wallis ANOVA (13/06/2025 16:00:08)

#### Notes

| | |
|---|---|
| X-Function | Kruskal-Wallis ANOVA |
| User Name | fgaro |
| Time | 13/06/2025 16:00:08 |
| Data Filter | No |

#### Input Data

| | Data | Range |
|---|---|---|
| Group Range | [Celegans]Sheet1!B | [1*:343*] |
| Data Range | [Celegans]Sheet1!C | [1*:343*] |

#### Descriptive Statistics

| | | N | Min | Q1 | Median | Q3 | Max |
|---|---|---|---|---|---|---|---|
| C | N2L1 | 33 | 440 | 879 | 882 | 884 | 887 |
| | N2L4 | 135 | 440 | 879 | 882 | 884 | 887 |
| | aex-3L1 | 12 | 440 | 879,75 | 883 | 884,75 | 885 |
| | aex-3L4 | 162 | 216 | 219 | 221,5 | 223 | 226 |

#### Ranks

| | | N | Mean Rank | Sum Rank |
|---|---|---|---|---|
| C | N2L1 | 33 | 245,57576 | 8104 |
| | N2L4 | 135 | 253,85185 | 34270 |
| | aex-3L1 | 12 | 256,33333 | 3076 |
| | aex-3L4 | 162 | 81,5 | 13203 |

#### Test Statistics

| | Chi-Square | DF | Prob>Chi-Square |
|---|---|---|---|
| C | 256,26436 | 3 | <0.0001 |

Null Hypothesis:The samples come from the same population.
Alternative Hypothesis:The samples come from different populations.
**C: At the 0.05 level, the populations are significantly different.**

#### Dunn's Test

| | | | Mean Rank Diff | Z | Prob | Sig |
|---|---|---|---|---|---|---|
| C | N2L1 | N2L4 | -8,27609 | -0,4319 | 1 | 0 |
| | N2L1 | aex-3L1 | -10,75758 | -0,3234 | 1 | 0 |
| | N2L1 | aex-3L4 | 164,07576 | 8,70621 | <0.0001 | 1 |
| | N2L4 | aex-3L1 | -2,48148 | -0,08348 | 1 | 0 |
| | N2L4 | aex-3L4 | 172,35185 | 14,9882 | <0.0001 | 1 |
| | aex-3L1 | aex-3L4 | 174,83333 | 5,92223 | <0.0001 | 1 |

Sig equals 1 indicates that the difference of the means is significant at the 0,05 level.
Sig equals 0 indicates that the difference of the means is NOT significant at the 0,05 level.

#### Paired Comparison Plot

C

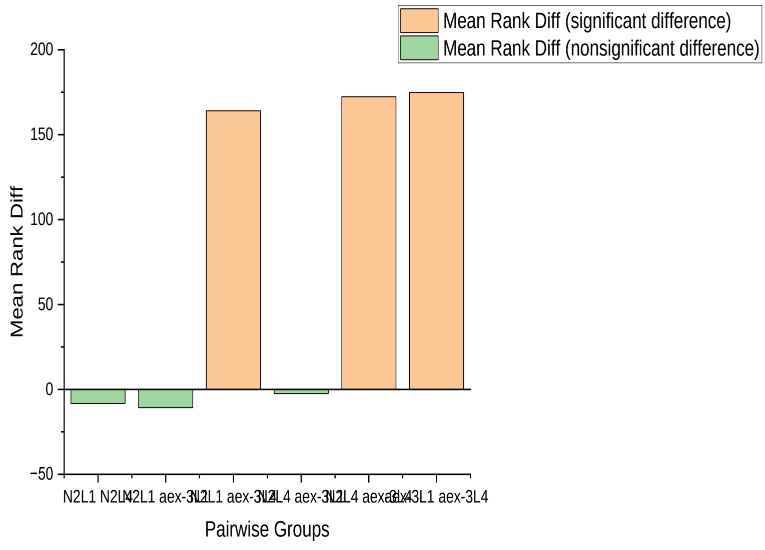
