## Supplementary material for "Tracking of *in vivo* O-GlcNAcylation in an Alzheimer’s Disease and Aging *C. elegans* Model": post_hoc_Analysis(Perim).pdf

### Kruskal-Wallis ANOVA (19/06/2025 16:31:30)

#### Notes

| | |
|---|---|
| X-Function | Kruskal-Wallis ANOVA |
| User Name | fgaro |
| Time | 19/06/2025 16:31:30 |
| Data Filter | No |

#### Input Data

| | Data | Range |
|---|---|---|
| Group Range | [Celegans]Sheet1!B | [1*:343*] |
| Data Range | [Celegans]Sheet1!D | [1*:343*] |

#### Descriptive Statistics

| | | N | Min | Q1 | Median | Q3 | Max |
|---|---|---|---|---|---|---|---|
| D | N2L1 | 33 | 105,32 | 105,32 | 105,32 | 105,32 | 105,32 |
| | N2L4 | 135 | 105,32 | 105,32 | 105,32 | 105,32 | 105,32 |
| | aex-3L1 | 12 | 105,32 | 105,32 | 105,32 | 105,32 | 105,32 |
| | aex-3L4 | 162 | 52,75 | 52,75 | 52,75 | 52,751 | 52,751 |

#### Ranks

| | | N | Mean Rank | Sum Rank |
|---|---|---|---|---|
| D | N2L1 | 33 | 252,5 | 8332,5 |
| | N2L4 | 135 | 252,5 | 34087,5 |
| | aex-3L1 | 12 | 252,5 | 3030 |
| | aex-3L4 | 162 | 81,5 | 13203 |

#### Test Statistics

| | Chi-Square | DF | Prob>Chi-Square |
|---|---|---|---|
| D | 313,16327 | 3 | <0.0001 |

Null Hypothesis:The samples come from the same population.
Alternative Hypothesis:The samples come from different populations.

**D: At the 0.05 level, the populations are significantly different.**

#### Dunn's Test

| | | | Mean Rank Diff | Z | Prob | Sig |
|---|---|---|---|---|---|---|
| D | N2L1 | N2L4 | 0 | 0 | 1 | 0 |
| | N2L1 | aex-3L1 | 0 | 0 | 1 | 0 |
| | N2L1 | aex-3L4 | 171 | 9,2415 | <0.0001 | 1 |
| | N2L4 | aex-3L1 | 0 | 0 | 1 | 0 |
| | N2L4 | aex-3L4 | 171 | 15,14577 | <0.0001 | 1 |
| | aex-3L1 | aex-3L4 | 171 | 5,89955 | <0.0001 | 1 |

Sig equals 1 indicates that the difference of the means is significant at the 0,05 level.
Sig equals 0 indicates that the difference of the means is NOT significant at the 0,05 level.

#### Paired Comparison Plot

D

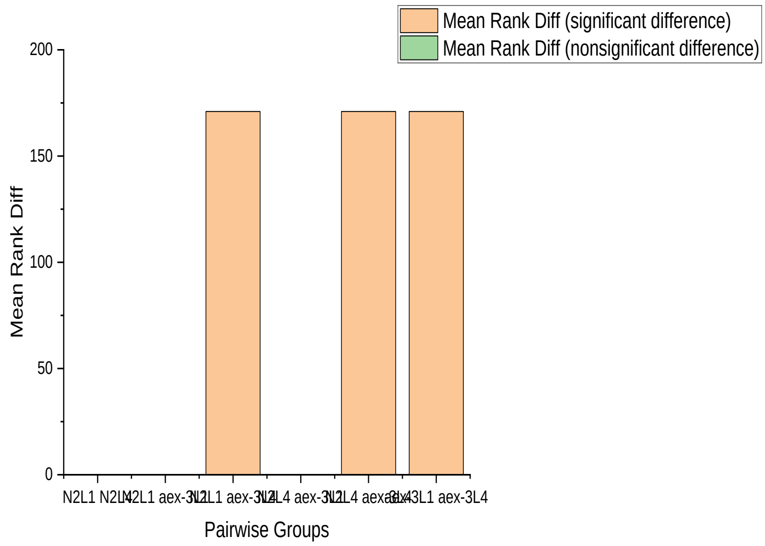
