## Supplementary material for "Tracking of *in vivo* O-GlcNAcylation in an Alzheimer’s Disease and Aging *C. elegans* Model": post_hoc_Analysis(x-traject).pdf

### Kruskal-Wallis ANOVA (25/06/2025 13:48:40)

#### Notes

| | |
|---|---|
| X-Function | Kruskal-Wallis ANOVA |
| User Name | fgaro |
| Time | 25/06/2025 13:48:40 |
| Data Filter | No |

#### Input Data

| | Data | Range |
|---|---|---|
| Group Range | [Celegans]Sheet1!B | [1*:343*] |
| Data Range | [Celegans]Sheet1!E | [1*:343*] |

#### Descriptive Statistics

| | | N | Min | Q1 | Median | Q3 | Max |
|---|---|---|---|---|---|---|---|
| E | N2L1 | 33 | 706 | 774,5 | 875 | 984 | 1663 |
| | N2L4 | 135 | 289 | 808 | 951 | 1298 | 1902 |
| | aex-3L1 | 12 | 710 | 752 | 820,5 | 922 | 1021 |
| | aex-3L4 | 162 | 798 | 923 | 1045 | 1094 | 1910 |

#### Ranks

| | | N | Mean Rank | Sum Rank |
|---|---|---|---|---|
| E | N2L1 | 33 | 117,09091 | 3864 |
| | N2L4 | 135 | 163,96667 | 22135,5 |
| | aex-3L1 | 12 | 81,20833 | 974,5 |
| | aex-3L4 | 162 | 195,54938 | 31679 |

#### Test Statistics

| | Chi-Square | DF | Prob>Chi-Square |
|---|---|---|---|
| E | 30,37292 | 3 | <0.0001 |

Null Hypothesis:The samples come from the same population.
Alternative Hypothesis:The samples come from different populations.
**E: At the 0.05 level, the populations are significantly different.**

#### Dunn's Test

| | | | Mean Rank Diff | Z | Prob | Sig |
|---|---|---|---|---|---|---|
| E | N2L1 | N2L4 | -46,87576 | -2,44158 | 0,08774 | 0 |
| | N2L1 | aex-3L1 | 35,88258 | 1,07666 | 1 | 0 |
| | N2L1 | aex-3L4 | -78,45847 | -4,15518 | 1,95021E-4 | 1 |
| | N2L4 | aex-3L1 | 82,75833 | 2,77884 | 0,03273 | 1 |
| | N2L4 | aex-3L4 | -31,58272 | -2,74125 | 0,03672 | 1 |
| | aex-3L1 | aex-3L4 | -114,34105 | -3,8657 | 6,64624E-4 | 1 |

Sig equals 1 indicates that the difference of the means is significant at the 0,05 level.
Sig equals 0 indicates that the difference of the means is NOT significant at the 0,05 level.

#### Paired Comparison Plot

E

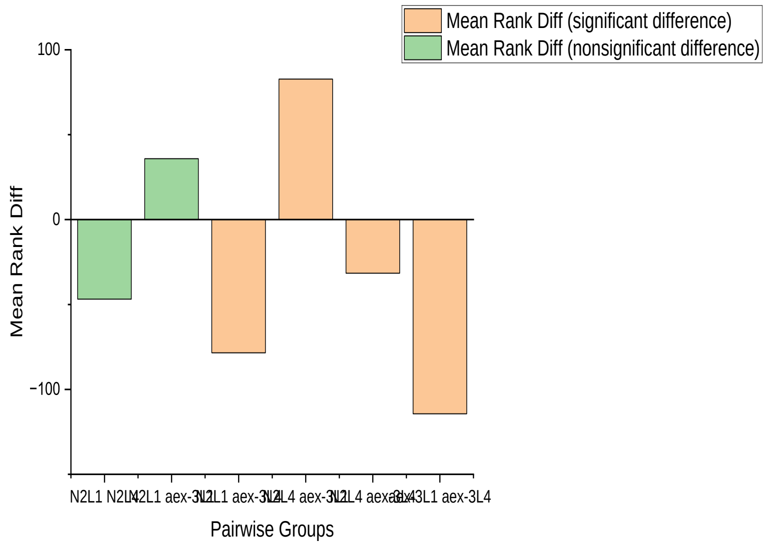
