## Supplementary material for "Tracking of *in vivo* O-GlcNAcylation in an Alzheimer’s Disease and Aging *C. elegans* Model": posthoc_y-trajectories.pdf

### Kruskal-Wallis ANOVA (25/06/2025 14:20:43)

#### Notes

| | |
|---|---|
| X-Function | Kruskal-Wallis ANOVA |
| User Name | fgaro |
| Time | 25/06/2025 14:20:43 |
| Data Filter | No |

#### Input Data

| | Data | Range |
|---|---|---|
| Group Range | [Celegans]Sheet1!B | [1*:343*] |
| Data Range | [Celegans]Sheet1!F | [1*:343*] |

#### Descriptive Statistics

| | | N | Min | Q1 | Median | Q3 | Max |
|---|---|---|---|---|---|---|---|
| F | N2L1 | 33 | 0 | 476 | 522 | 559,5 | 906 |
| | N2L4 | 135 | 0 | 345 | 499 | 644 | 1106 |
| | aex-3L1 | 12 | 0 | 464,5 | 535 | 567,75 | 656 |
| | aex-3L4 | 162 | 441 | 581 | 645 | 701 | 1059 |

#### Ranks

| | | N | Mean Rank | Sum Rank |
|---|---|---|---|---|
| F | N2L1 | 33 | 127,72727 | 4215 |
| | N2L4 | 135 | 131,98889 | 17818,5 |
| | aex-3L1 | 12 | 116,125 | 1393,5 |
| | aex-3L4 | 162 | 217,44444 | 35226 |

#### Test Statistics

| | Chi-Square | DF | Prob>Chi-Square |
|---|---|---|---|
| F | 66,77901 | 3 | <0.0001 |

Null Hypothesis:The samples come from the same population.
Alternative Hypothesis:The samples come from different populations.
**F: At the 0.05 level, the populations are significantly different.**

#### Dunn's Test

| | | | Mean Rank Diff | Z | Prob | Sig |
|---|---|---|---|---|---|---|
| F | N2L1 | N2L4 | -4,26162 | -0,22197 | 1 | 0 |
| | N2L1 | aex-3L1 | 11,60227 | 0,34812 | 1 | 0 |
| | N2L1 | aex-3L4 | -89,71717 | -4,7514 | <0.0001 | 1 |
| | N2L4 | aex-3L1 | 15,86389 | 0,53267 | 1 | 0 |
| | N2L4 | aex-3L4 | -85,45556 | -7,41712 | <0.0001 | 1 |
| | aex-3L1 | aex-3L4 | -101,31944 | -3,42543 | 0,00368 | 1 |

Sig equals 1 indicates that the difference of the means is significant at the 0,05 level.
Sig equals 0 indicates that the difference of the means is NOT significant at the 0,05 level.

#### Paired Comparison Plot

F

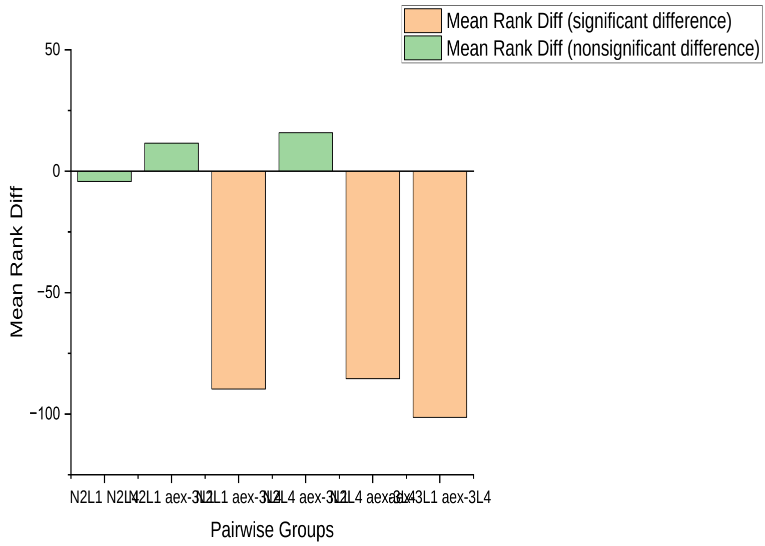
