## Supplementary figures and images for "Tracking of *in vivo* O-GlcNAcylation in an Alzheimer’s Disease and Aging *C. elegans* Model"

### 64x64.png

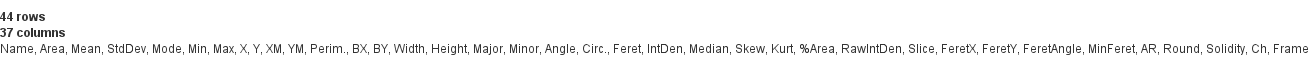

### 64x64.png

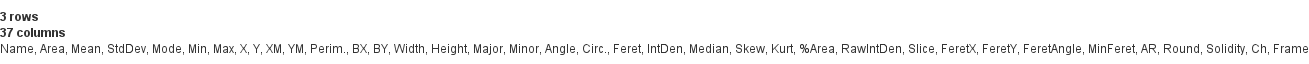

### 64x64.png

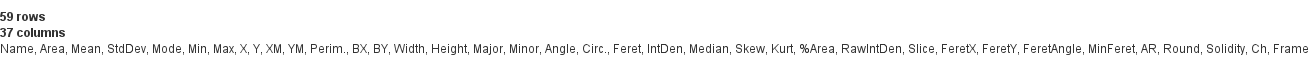

### 64x64.png

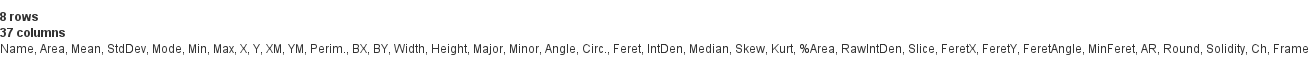

### 128x128.png

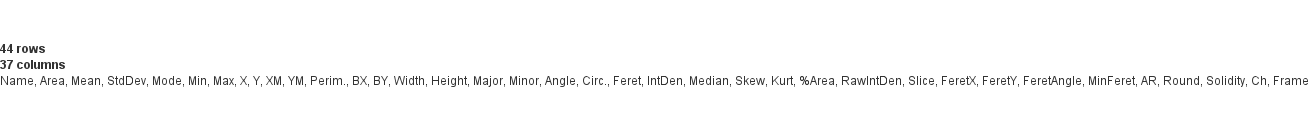

### 128x128.png

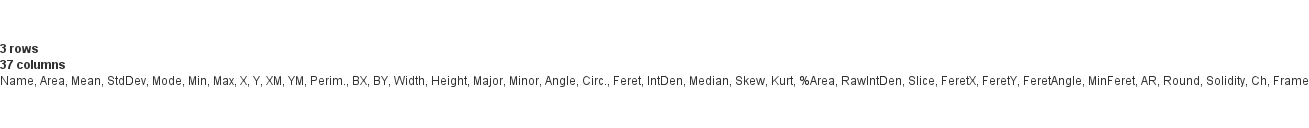

### 128x128.png

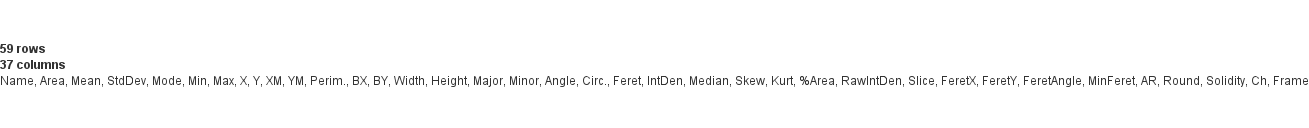

### 128x128.png

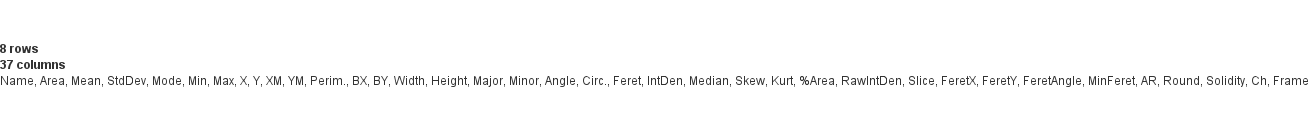

### 256x256.png

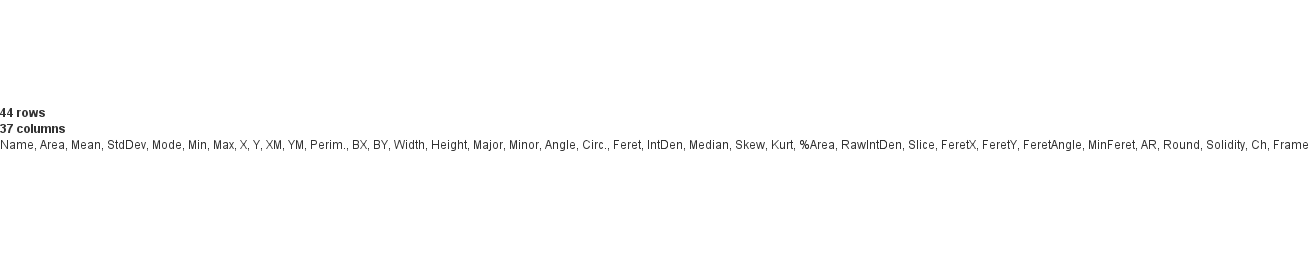

### 256x256.png

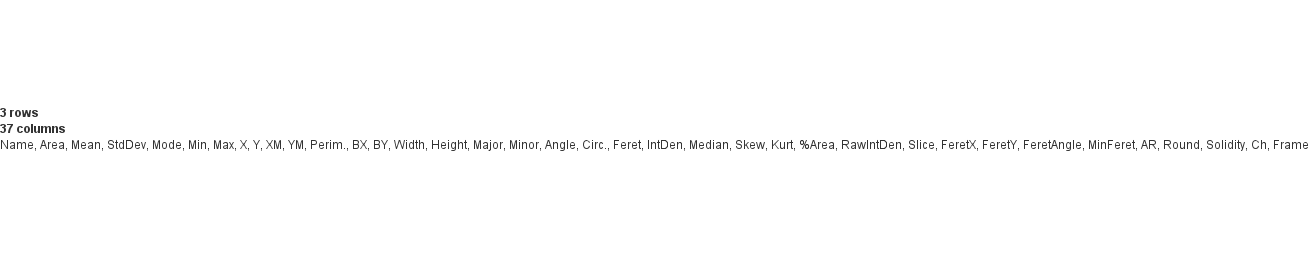

### 256x256.png

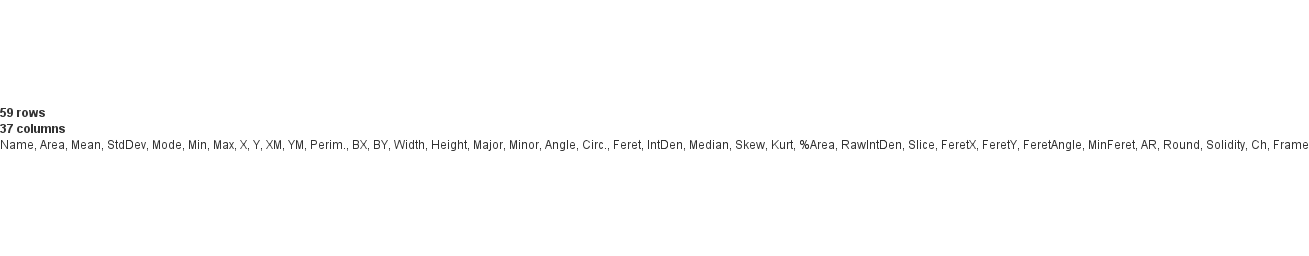

### 256x256.png

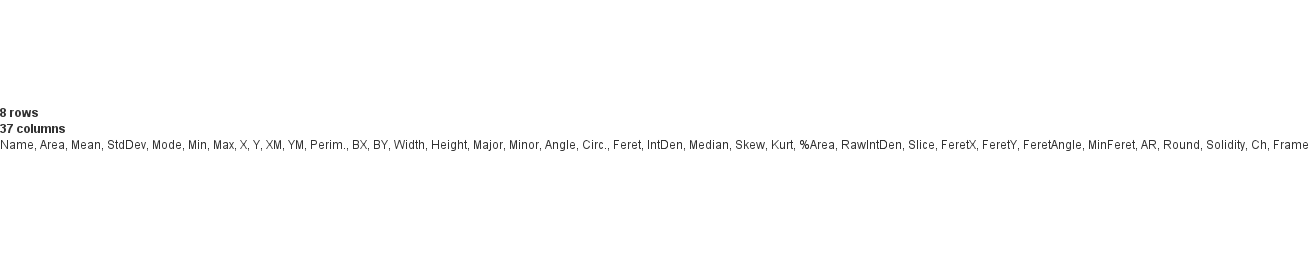
